# Structural basis of actin filament assembly and aging

**DOI:** 10.1101/2022.03.29.486216

**Authors:** Wout Oosterheert, Björn U Klink, Alexander Belyy, Sabrina Pospich, Stefan Raunser

**Author notes:** Correspondence to Stefan Raunser.

## Abstract

The dynamic turnover of actin filaments (F-actin) controls cellular motility in eukaryotes and is coupled to changes in the F-actin nucleotide state. It remains unclear how F-actin hydrolyzes ATP and subsequently undergoes subtle conformational rearrangements that ultimately lead to filament depolymerization by actin-binding proteins. Here, we present cryo-EM structures of F-actin in all nucleotide states, polymerized in the presence of Mg^2+^ or Ca^2+^, at resolutions (∼2.2 Å) that allow for the visualization of hundreds of water molecules. The structures reveal that the G- to F-actin transition induces the relocation of water molecules in the nucleotide binding pocket, activating one of them for the nucleophilic attack of ATP. Unexpectedly, the back door for the subsequent release of inorganic phosphate (P_i_) is closed in all structures, indicating that the F-actin conformation that allows for P_i_ release occurs transiently. The small changes in the nucleotide-binding pocket after ATP hydrolysis and P_i_ release are sensed by a key amino acid, amplified and transmitted to the filament periphery. Furthermore, differences in the positions of waters in the nucleotide binding pocket explain why Ca^2+^-actin exhibits slower polymerization rates than Mg^2+^-actin. Our work elucidates the solvent-driven rearrangements that govern actin filament assembly and aging and lays the foundation for the rational design of drugs and small molecules for imaging and therapeutic applications.

## Introduction

The cytoskeletal protein actin transitions between monomeric, globular (G-actin) and polymeric, filamentous (F-actin) states to control the shape, polarity and migration of eukaryotic cells^1–3^. Many processes driven by actin, such as cell division, depend on its ATPase activity^1^; ATP binds with nanomolar affinity in a central pocket in the enzyme, but actin exhibits only very weak ATPase activity (7 10^-6^ s^-1^) in its monomeric form^4^. Polymerization triggers a conformational change in the nucleotide binding pocket that allows actin to hydrolyze ATP within seconds (0.3 s^-1^) of filament formation^5^. The cleaved inorganic phosphate (P_i_) is not released immediately after hydrolysis (release rate 0.006 s^-1^)^6^, yielding the metastable intermediate ADP-P_i_ state of F-actin^7^. After the exit of P_i_, ADP-bound F-actin represents the ‘aged’ state of the filament, which can then be depolymerized back to G-actin. *In vivo*, this cyclic process is tightly regulated by a wide variety of actin-binding proteins (ABPs) that control the kinetics of actin turnover^8–10^. For instance, G-actin probably does not exist at significant concentrations as the uncomplexed monomer *in vivo*; instead it is essentially always bound to ABPs such as profilin and thymosin-*β*4 to prevent uncontrolled nucleation events^11, 12^. Interestingly, a subset of ABPs is capable of directly sensing the nucleotide state of either G- or F-actin^13, 14^. As prominent example, ABPs of the ADF/cofilin family efficiently bind and sever F-actin in the ADP state to promote actin turnover, but bind only with weaker affinity to ‘young’ actin filaments that harbor ATP or ADP-P_i_ in their active site^15–17^. In addition to ABPs, the divalent cation that associates with the actin-bound nucleotide, magnesium (Mg^2+^) or calcium (Ca^2+^), also strongly affects polymerization rates. Both Mg^2+^ and Ca^2+^ bind to ATP-actin with nanomolar affinity^18, 19^, but polymerization rates of Mg^2+^-actin are much faster than those of Ca^2+^-actin^18, 20, 21^. Determining whether Mg^2+^ or Ca^2+^ associates with the actin nucleotide *in vivo* was a major focus in the early days of actin research. A 1969 study proposed that Mg^2+^ is the cation bound to actin in native myofibrils^22^, because EDTA-treatment could not deplete Mg^2+^ levels in the myofibril preparations. Combined with further evidence from electron probe analyses of the skeletal muscle I-band^23^ and the observation that the Mg^2+^ concentration in most cell types is orders of magnitude higher than the Ca^2+^ concentration^24^, it is now accepted that Mg^2+^ is the cation that predominantly associates with the nucleotide of actin *in vivo*^25^. Although Ca^2+^ probably binds only marginally to the actin-nucleotide in a physiological setting, it is standardly used in actin purification protocols^26, 27^. The Ca^2+^-ATP bound form of G-actin exhibits slower polymerization kinetics and a higher critical concentration of polymerization^25, 28^, making it easier to retain actin in its G-form during purification procedures for *in vitro* studies. Accordingly, ∼70% of all G-actin crystal structures were solved with Ca^2+^ bound in the active site^29^. Comparing these crystal structures to those solved with Mg^2+^ reveals that the coordination shells of Ca^2+^ and Mg^2+^ differ when bound to the nucleotide of G-actin, with a typical hepta-coordinated, pentagonal bipyramidal arrangement for Ca^2+^ and a hexa-coordinated, octahedral arrangement for Mg^2+^. Despite this difference in cation-coordination, the nucleotide-binding sites and overall conformations of Ca^2+^-G-actin and Mg^2+^-G-actin are essentially the same^30^, suggesting that rearrangements in the filamentous state cause the slow polymerization rates of Ca^2+^-actin.

While a large number of G-actin structures have been solved by x-ray crystallography since 1990^30, 31^, structural models of F-actin have only been obtained by other methods because actin filaments resist crystallization. The actin filament can be described as a double-stranded, right handed helix^32^. A fiber diffraction-derived model of F-actin was reported in 2009 and revealed that the actin monomer flattens upon incorporation into the filament^33^. In 2015, our group published the 3.7-Å single-particle cryo-EM structure of F-actin in complex with the ABP tropomyosin^34^, which represented the first complete atomic model of actin in the filamentous state. Since then, numerous cryo-EM studies by our group and others have revealed the F-actin architecture in all nucleotide states^35, 36^ and in complex with a variety of ABPs such as cofilin^37, 38^ and myosin^39–41^. However, all previously published F-actin structures were solved at moderate resolutions of ∼3 - 4.5 Å, and therefore did not display sufficient details to allow for the modeling of solvent molecules and exact positions of amino-acid side chains. Hence, key mechanistic events in the aging process of F-actin such as ATP hydrolysis, which strongly depends on water molecules, are still not understood. In addition, the currently available structural data do not reveal how changes in the nucleotide state in the F-actin interior are transmitted to the surface of the filament where they can be sensed by ABPs, nor do they explain the differences in polymerization kinetics between Ca^2+^-bound and Mg^2+^-bound actin. Here, we present ∼2.2 Å cryo-EM structures of rabbit skeletal *α*-actin filaments in all three functional states polymerized in the presence of Mg^2+^ or Ca^2+^. The structures illuminate the F-actin architecture in unprecedented detail and underpin the critical role of solvent molecules in actin filament assembly and aging.

## Results and Discussion

### Cryo-EM structures of F-actin at 2.2 Å resolution allow for solvent modeling

To structurally understand the turnover of F-actin that goes along with ATP hydrolysis and slight conformational changes that can be sensed by actin-binding proteins, structures of F-actin are needed at a resolution that allows for the reliable modelling of side chains, the nucleotide, phosphate, metal ions and water molecules. Therefore, we developed a cryo-EM data collection and image processing workflow to greatly improve the resolution of previous F-actin reconstructions (Supplementary Fig. 1, Methods). By employing this optimized workflow, we determined structures of Mg^2+^-F-actin in all relevant nucleotide states (ATP, ADP-P_i_, ADP) at resolutions of 2.17 – 2.24 Å, which is suitable for the modeling of solvent molecules (Fig. 1a-d, Supplementary Fig. 2, 4, Table 1, Methods).

**Fig. 1.**
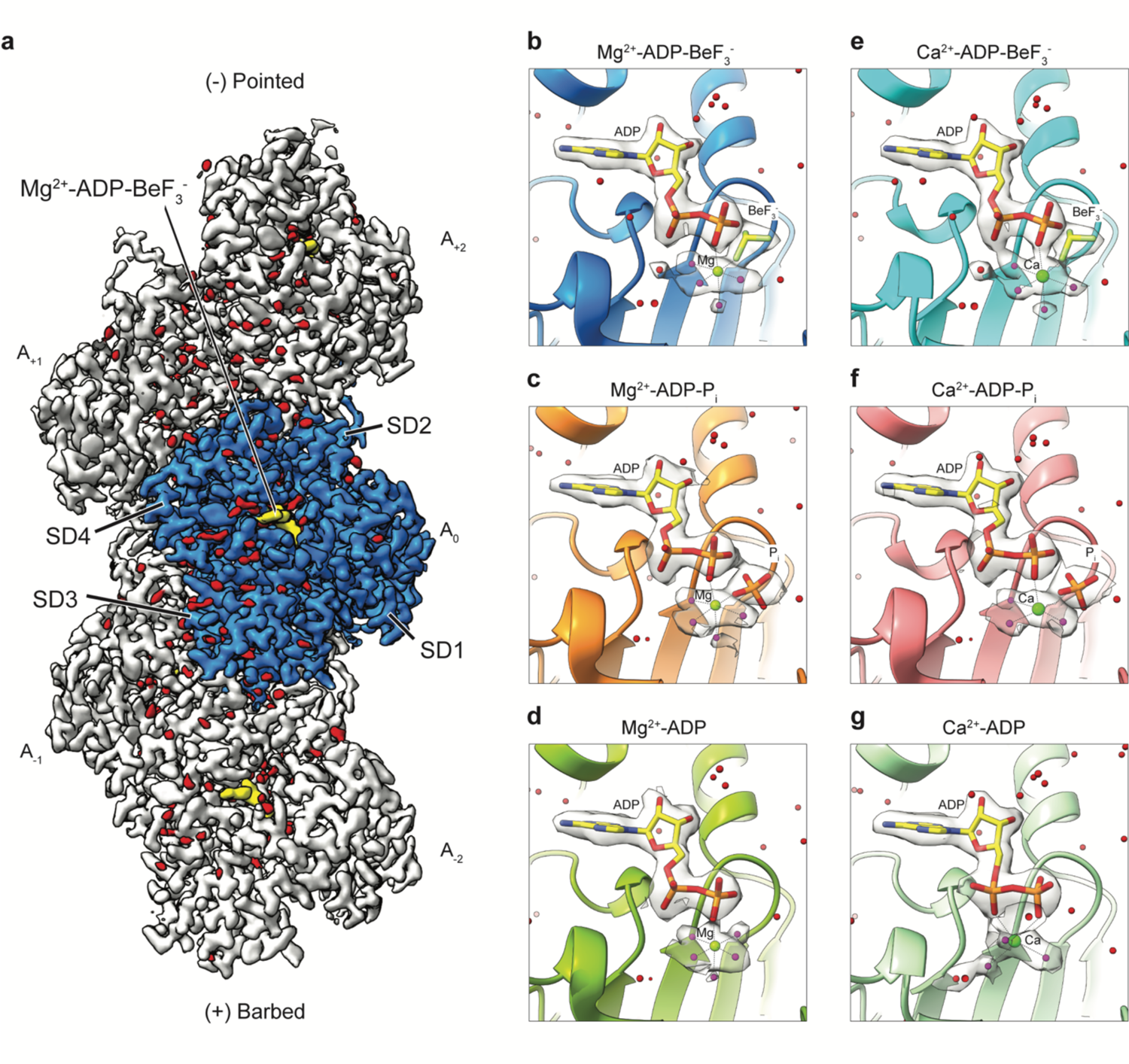
Cryo-EM reconstructions of F-actin at 2.2-Å resolution. **a**, Local-resolution filtered, sharpened cryo-EM density map of F-actin in the Mg^2+^-ADP-BeF_3_^-^ state. The subunits are labeled based on their location along the filament, ranging from the barbed (A_-2_) to the pointed (A_2_) end. The central actin subunit (A_0_) is colored blue, the other four subunits are grey. Actin subdomains (SD1-4) are annotated in the central subunit. Densities corresponding to water molecules are colored red. **b-g** Cryo-EM densities of the nucleotide-binding pocket in F-actin in the Mg^2+^-ADP-BeF_3_^-^ **(b)**, Mg^2+^-ADP-P_i_ **(c)**, Mg^2+^-ADP **(d)**, Ca^2+^-ADP-BeF_3_^-^ **(e)**, Ca^2+^-ADP-P_i_ **(f)** and Ca^2+^-ADP **(g)** states. Waters that directly coordinate the nucleotide-associated cation are colored magenta. For the Ca^2+^-ADP structure **(g)**, one coordinating water is hidden behind the Ca^2+^-ion.

**Table 1.** Cryo-EM data collection and processing of the Mg^2+^-F-actin datasets

| F-actin state | Mg<sup>2+</sup>-ADP-BeF<sub>3</sub><sup>-</sup><br>PDB XXXX<br>EMD-XXXX | Mg<sup>2+</sup>-ADP-P<sub>i</sub><br>PDB XXXX<br>EMD-XXXX | Mg<sup>2+</sup>-ADP<br>PDB XXXX<br>EMD-XXXX |
| --- | --- | --- | --- |
| **Data collection and processing** |  |  |  |
| Microscope | Titan Krios | Titan Krios | Titan Krios |
| Camera | K3 | K3 | K3 |
| Magnification | 130,000 | 130,000 | 130,000 |
| Voltage (kV) | 300 | 300 | 300 |
| Exposure time frame/total (s) | 0.05/3 | 0.0375/3 | 0.05/3 |
| Number of frames | 60 | 80 | 60 |
| Electron exposure (e<sup>-</sup>/Å<sup>2</sup>) | 78.9 | 88.8 | 76.4 |
| Defocus range (μm) | -0.7 to -2.0 | -0.7 to -2.0 | -0.7 to -2.0 |
| Pixel size (Å) | 0.695 | 0.695 | 0.695 |
| Micrographs (no.) | 10,822 | 9,658 | 9,842 |
| Initial particle images (no.) | 2,897,679 | 2,349,979 | 1,296,776 |
| Final particles images (no.) | 2,228,553 | 1,808,554 | 1,114,051 |
| Map Resolution | 2.17 | 2.22 | 2.24 |
| 0.143 FSC threshold |  |  |  |
| Map resolution range (Å) | 2.1 – 3.2 | 2.2 – 3.2 | 2.2 – 3.4 |
| Measured helical symmetry<sup>a</sup> |  |  |  |
| Helical rise (Å) | 27.47±0.05 | 27.50±0.02 | 27.50±0.07 |
| Helical twist (°) | 166.6±0.2 | 166.5±0.4 | 166.7±0.1 |
| **Refinement** |  |  |  |
| Model Resolution (Å) | 2.20 | 2.28 | 2.35 |
| 0.5 FSC threshold |  |  |  |
| Map sharpening B factor (Å<sup>2</sup>) | -50 | -64 | -65 |
| Model composition |  |  |  |
| Non-hydrogen atoms | 14,990 | 15,141 | 15,046 |
| Protein residues | 1,830 | 1,835 | 1,830 |
| Waters | 480 | 591 | 556 |
| Ligands | 15 | 15 | 10 |
| B factors (Å<sup>2</sup>) |  |  |  |
| Protein | 31.53 | 34.56 | 38.07 |
| Waters | 31.82 | 23.19 | 31.98 |
| Ligands | 22.89 | 31.58 | 31.71 |
| R.m.s. deviations |  |  |  |
| Bond lengths (Å) | 0.004 | 0.008 | 0.003 |
| Bond angles (°) | 0.592 | 0.669 | 0.549 |
| Validation |  |  |  |
| EM-ringer score | 5.94 | 6.45 | 5.89 |
| MolProbity score | 1.17 | 1.12 | 1.03 |
| Clashscore | 2.75 | 3.33 | 2.50 |
| Rotamers outliers (%) | 0.65 | 0.64 | 0.32 |
| Ramachandran plot |  |  |  |
| Favored (%) | 97.5 | 98.1 | 98.05 |
| Allowed (%) | 2.5 | 1.9 | 1.95 |
| Outliers (%) | 0 | 0 | 0 |

<sup>a</sup>The helical parameters were estimated from the atomic model of five consecutive subunits independently fitted to the map as described previously (Pospich et al., 2017).

The ATP state of F-actin is short lived, which required the use of an ATP analog for our structural studies. We considered that previous work from our group has shown that the widely used non-hydrolysable ATP analog AppNHp (also known as AMP-PNP) is a suboptimal ligand for F-actin because its degradation product, the ADP analog AppNH_2_, exhibits higher affinity for F-actin^42^ and hence accumulates in the active site during filament preparation^35^. We therefore opted to determine F-actin structures using ADP complexed with beryllium fluoride (BeF_3_^-^, also referred to as BeF_x_)^43^ to mimic the ATP state of the actin filament (Fig. 1b).

The unprecedentedly high resolutions of the F-actin reconstructions allowed for the modelling of hundreds of solvent molecules, i.e. waters, and for the precise positioning of the majority of amino-acid side chains in each density map (Fig. 1, Supplementary Fig. 4, 7). Accordingly, we observed a clear density for the nucleotide, the associated divalent cation Mg^2+^ and the cation-coordinating waters (Fig. 1b-d, Movie S1). The overall structures of all nucleotide states of Mg^2+^-F-actin are highly similar, with a C*α* atom root-mean square deviation (r.m.s.d.) of < 0.6 Å (Supplementary Fig. 6a), indicating that the differences are in the details (see below). In addition, F-actin in all three structures has the same helical rise and twist (Table 1). These results are in line with the previous findings that the stable states adopted by F-actin during its aging process share identical helical parameters^35, 36^. Surprisingly, although earlier studies predicted additional Mg^2+^ and P_i_ binding sites outside of the F-actin nucleotide-binding pocket^29, 44, 45^, we did not find evidence for these secondary ion-binding sites in any of our reconstructions.

In the Mg^2+^-ADP-BeF_3_^-^ F-actin structure (2.17 Å), we observed unambiguous density for the modeling of BeF_3_^-^ in the nucleotide-binding site (Fig. 1b). The nucleotide conformation in Mg^2+^-ADP-BeF_3_^-^ differed from Mg^2+^-ADP-P_i_, but instead resembled Mg^2+^-ATP in G-actin (Fig. 2b, c Supplementary Fig. 9a). Accordingly, the distance between the oxygen of the β-phosphate (P*_β_*) of ADP and Be (1.4 Å) is as short as the equivalent distance in ATP (1.5 Å), indicating that BeF_3_^-^ binds covalently to ADP as has been previously suggested^46^. P_i_ resides at much larger distance from ADP (2.9 Å) (Supplementary Fig. 9a). Thus, our structures rationalize why BeF_3_^-^ binds with a much higher affinity (∼1-2 μM)^43^ to ADP F-actin than P_i_ (∼1.5 mM)^47^. Previous reports suggested that actin filaments with ADP-BeF_3_^-^ incorporated in their active site adopt a conformation similar to the transition state ADP-P*^43^ or even the ADP-P_i_-like state of the filament^48, 49^. However, the Mg^2+^-ADP-BeF_3_^-^ F-actin structure reveals no transition state or ADP-P_i_-like arrangement of the nucleotide and instead defines ADP-BeF_3_^-^ as a mimic of the ATP ground state of F-actin, which is consistent with observations for other molecular systems^50, 51^.

**Fig. 2.**
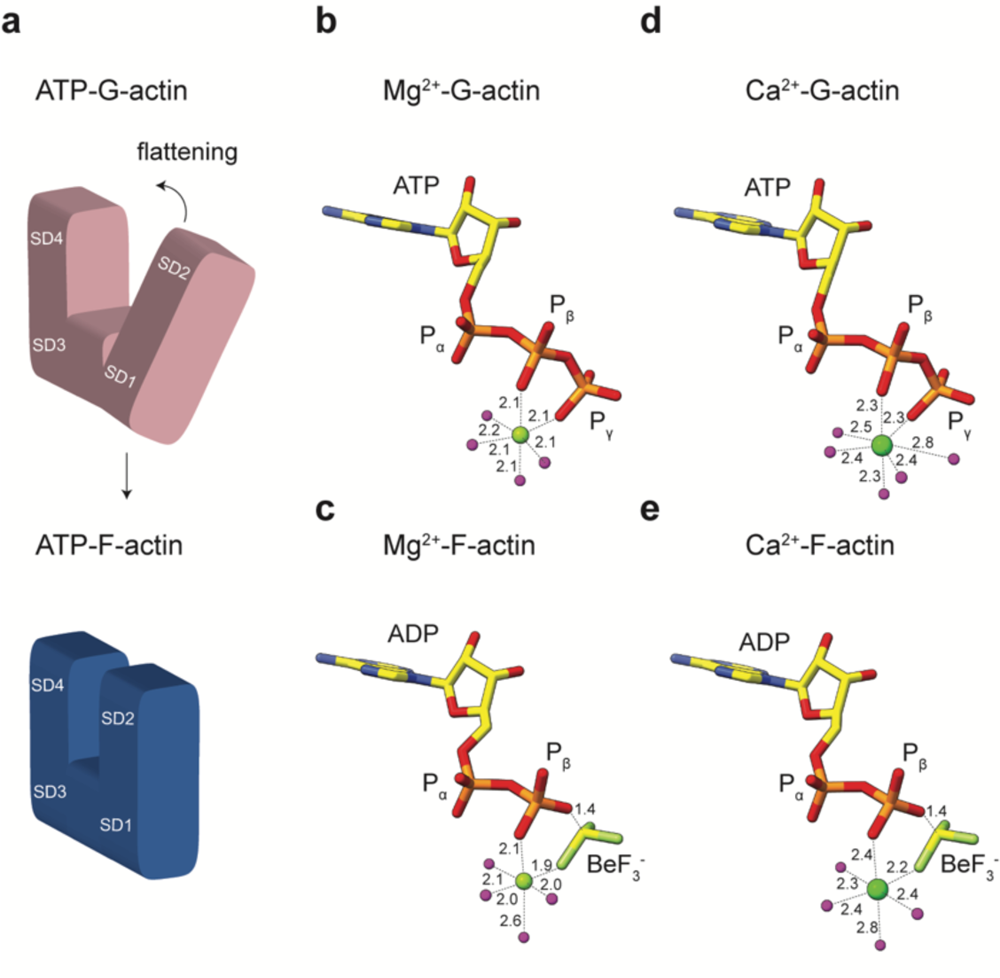
Cation coordination at the nucleotide-binding site of ATP-actin. **a** Schematic cartoon representation of actin flattening during the G- to F-actin transition. **b-e** Nucleotide conformation and inner-coordination sphere of the divalent cation in Mg^2+^-ATP G-actin (pdb 2v52) **(b)**, Mg^2+^-ADP-BeF_3_^-^ F-actin **(c)**, Ca^2+^-ATP G-actin (pdb 1qz5) **(d)**, and Ca^2+^-ADP-BeF_3_^-^ F-actin **(e)**. Bond lengths are annotated in Å.

In order to elucidate the structural and mechanistic basis for the slower polymerization kinetics of Ca^2+^-actin, we furthermore aimed to solve F-actin structures in all relevant nucleotide states with Ca^2+^ instead of Mg^2+^ as nucleotide-associated cation. To this end, we prepared actin filaments without the addition of EGTA and Mg^2+^ before polymerization (Methods), and obtained Ca^2+^-F-actin structures (resolution between brackets) in complex with ADP-BeF_3_^-^ (2.21 Å), ADP-P_i_ (2.15 Å) and ADP (2.15 Å) (Fig. 1e-g, Supplementary Fig. 3, 5, 8 Table 2, Movie S2, Methods). The high quality of the Ca^2+^-F-actin reconstructions revealed that, even though Ca^2+^-actin displays slow polymerization and fast depolymerization kinetics^18^, it adopts stable conformations in the filamentous state. Globally, the Ca^2+^-F-actin structures are comparable to those of Mg^2+^-F-actin, with no changes in helical rise and twist and a C*α* atom r.m.s.d. of < 0.6 Å (Supplementary Fig. 6b-e). Thus, contrary to previous studies that reported large conformational changes between Mg^2+^ and Ca^2+^-F-actin based on low-resolution cryo-EM reconstructions^52, 53^, our ∼2.2 Å structures indicate that the change of divalent cation from Mg^2+^ to Ca^2+^ does not induce any large conformational rearrangements in the filament.

**Table 2.** Cryo-EM data collection and processing of the Ca^2+^-F-actin datasets

| F-actin state | Ca<sup>2+</sup>-ADP-BeF<sub>3</sub><sup>-</sup><br>PDB XXXX<br>EMD-XXXX | Ca<sup>2+</sup>-ADP-P<sub>i</sub><br>PDB XXXX<br>EMD-XXXX | Ca<sup>2+</sup>-ADP<br>PDB XXXX<br>EMD-XXXX |
| --- | --- | --- | --- |
| **Data collection and processing** |  |  |  |
| Microscope | Titan Krios | Titan Krios | Titan Krios |
| Camera | K3 | K3 | K3 |
| Magnification | 130,000 | 130,000 | 130,000 |
| Voltage (kV) | 300 | 300 | 300 |
| Exposure time frame/total (s) | 0.05/3 | 0.0375/3 | 0.05/3 |
| Number of frames | 60 | 80 | 60 |
| Electron exposure (e<sup>-</sup>/Å<sup>2</sup>) | 84.6 | 88.8 | 75.6 |
| Defocus range (μm) | -0.7 to -2.0 | -0.7 to -2.0 | -0.7 to -2.0 |
| Pixel size (Å) | 0.695 | 0.695 | 0.695 |
| Micrographs (no.) | 10,705 | 10,156 | 10,733 |
| Initial particle images (no.) | 2,246,622 | 3,031,270 | 1,873,773 |
| Final particles images (no.) | 1,719,432 | 2,171,987 | 1,073,455 |
| Map Resolution | 2.20 | 2.15 | 2.15 |
| 0.143 FSC threshold |  |  |  |
| Map resolution range (Å) | 2.15 – 3.2 | 2.1 – 3.1 | 2.1 – 3.5 |
| Measured helical symmetry<sup>a</sup> |  |  |  |
| Helical rise (Å) | 27.46±0.05 | 27.47±0.08 | 27.47±0.05 |
| Helical twist (°) | 166.7±0.1 | 166.6±0.1 | 166.5±0.1 |
| **Refinement** |  |  |  |
| Model Resolution (Å) | 2.23 | 2.16 | 2.17 |
| 0.5 FSC threshold |  |  |  |
| Map sharpening B factor (Å<sup>2</sup>) | -55 | -55 | -55 |
| Model composition |  |  |  |
| Non-hydrogen atoms | 15,022 | 15,078 | 15,060 |
| Protein residues | 1,830 | 1,835 | 1,830 |
| Waters | 512 | 528 | 570 |
| Ligands | 15 | 15 | 10 |
| B factors (Å<sup>2</sup>) |  |  |  |
| Protein | 24.67 | 21.12 | 26.49 |
| Waters | 23.66 | 23.30 | 28.02 |
| Ligands | 16.14 | 15.86 | 20.53 |
| R.m.s. deviations |  |  |  |
| Bond lengths (Å) | 0.003 | 0.004 | 0.003 |
| Bond angles (°) | 0.542 | 0.596 | 0.568 |
| Validation |  |  |  |
| EM-ringer score | 6.31 | 6.39 | 6.56 |
| MolProbity score | 1.17 | 1.02 | 0.99 |
| Clashscore | 2.75 | 2.39 | 1.81 |
| Rotamers outliers (%) | 0.65 | 0.64 | 0.00 |
| Ramachandran plot |  |  |  |
| Favored (%) | 97.5 | 98.1 | 97.8 |
| Allowed (%) | 2.5 | 1.9 | 1.9 |
| Outliers (%) | 0 | 0 | 0.3 |

<sup>a</sup>The helical parameters were estimated from the atomic model of five consecutive subunits independently fitted to the map as described previously (Pospich et al., 2017).

### The G- to F-actin transition triggers the relocation of water molecules and ATP hydrolysis

A comparison of previously reported high-resolution crystal structures of rabbit G-actin in the ATP state^54, 55^ with our high-resolution cryo-EM structures of the ATP state of F-actin allow, for the first time, a description of the G- to F-actin transition at an unprecedented level of molecular detail. Upon polymerization, subdomains 1 and 2 (SD1 and SD2) of the actin monomer rotate ∼15°, leading to a more compact arrangement in the filament (Fig. 2a). This process, in which the backbones of residues Q137 and K336 act as hinges (as determined by DynDom^56^) (Fig. 3a, b), is commonly referred to as actin monomer flattening^33^.

**Fig. 3.**
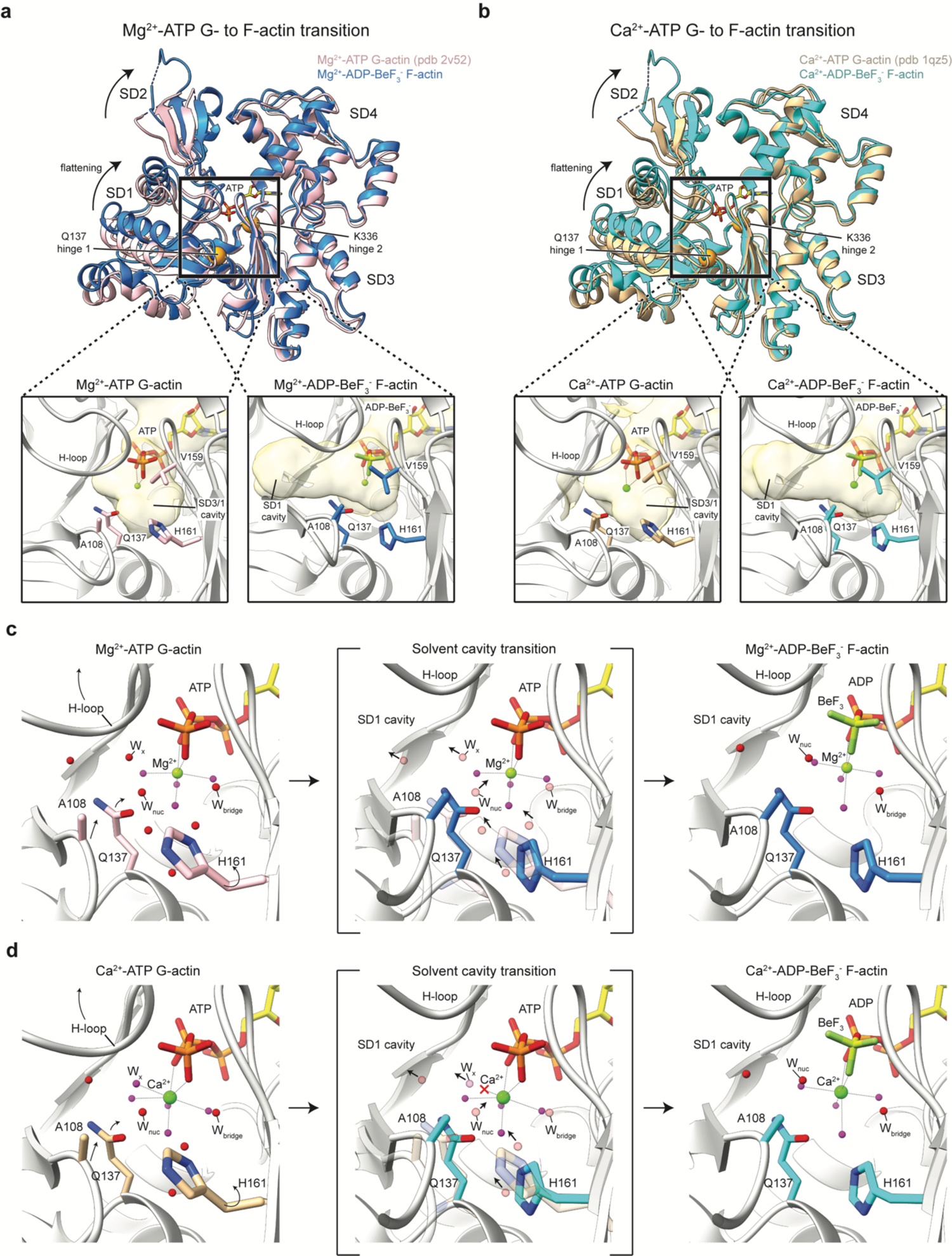
Solvent cavity remodeling during the G- to F-actin transition. **a, b** Upper panels: Overlay of G- and F-actin structures reveal the global conformational changes associated with actin flattening for Mg^2+^-**(a)** and Ca^2+^-F-actin **(b)**. Hinges Q137 and K336 are shown as orange spheres. Lower panels: Internal solvent cavities near the nucleotide-binding site in G- and F-actin. The cavities were calculated by the Castp3 server and are shown as beige, semi-transparent surfaces. The nucleotide was not considered in the solvent cavity calculations. **c, d** Water relocation in Mg^2+^-**(c)** and Ca^2+^-actin **(d)**. Waters that directly coordinate the nucleotide-associated cation are colored magenta. Left panel: water and amino-acid arrangement in ATP-G-actin. Amino acids are colored pink for Mg^2+^-actin and light-brown for Ca^2+^-actin, whereas the cartoon representation is shown in grey. Arrows depict the movement of amino-acid regions for the transition to F-actin. Middle panel: Overlay of the amino-acid positions in ADP-BeF_3_^-^ F-actin (colored blue for Mg^2+^-actin and cyan for Ca^2+^-actin) with the solvent molecules in the G-actin structure. The waters in the SD3/1 cavity of ATP-G-actin are shown as semi-transparent spheres. Arrows indicate the direction of water relocation. Right panel: Water and amino-acid arrangement in ADP-BeF_3_^-^ F-actin.

We first analyzed how actin flattening corresponds with the arrangement of the bound nucleotide. In Mg^2+^-ADP-BeF_3_^-^ F-actin, the nucleotide adopts a conformation comparable to ATP in G-actin (Fig. 1b, 2b, c). The associated Mg^2+^ ion is coordinated by P*_β_* of ADP, a fluoride moiety of BeF_3_^-^ and four water molecules, defining a hexa-coordinated, octahedral coordination, similar to that of Mg^2+^ in ATP-G-actin (Fig. 1b, 2b, c). Our F-actin structure thus provides strong experimental evidence that Mg^2+^ retains its water coordination during the G- to F-actin transition, which was previously only predicted based on molecular dynamics (MD) data^29^.

Since the high-resolution F-actin structures visualize solvent molecules, it is now possible to pinpoint the relocation of water molecules during the G- to F-actin transition. In Mg^2+^-ATP G-actin (pdb 2v52)^55^, there is a large cavity (∼7 Å diameter) that accommodates several ordered water molecules in front of the ATP *γ*-phosphate between SD3 and SD1 (SD3/1 cavity, Fig. 3a). Actin flattening results in the upward displacement of the H-loop (residues 72 – 77) and the movement of the proline-rich loop (residues 108 – 112) and the side chains of Q137 and H161 towards the nucleotide (Fig. 3c). As a result, the SD3/1 cavity becomes narrower (∼5 Å diameter) and a new cavity in the SD1 (deemed SD1 cavity) opens up (Fig. 3a). Due to the narrowing of the SD3/1 cavity, several waters in this cavity would clash with amino acids and therefore need to relocate to the SD1 cavity through a path that involves the movement of a water (W_x_) that is bound in between both cavities (Fig. 3c, Movie S3). Importantly, the relocation of water molecules into the SD1 cavity does not impact those coordinating the nucleotide-bound Mg^2+^ ion, and hence the Mg^2+^ coordination sphere does not change between G- and F-actin (Fig. 2b).

After the conformational change from G- to F-actin, only three water molecules remain in the SD3/1 cavity that are not coordinated by Mg^2+^. One of them is hydrogen-bonded to the C=O group of the Q137 side chain (Fig. 4a, Supplementary Fig. 10b). Due to the rearrangement of the nucleotide-binding site in F-actin, this water resides within much closer distance (3.6 Å) to the P*γ*-analog BeF_3_^-^ (Fig. 4a, Supplementary Fig. 10b) than in Mg^2+^-ATP G-actin (>4 Å distance from the P*γ* (4.6 Å in pdb 2v52, Supplementary Fig. 10a)^55^). Since no other ordered waters align in front of the nucleotide, the water that is hydrogen-bonded to Q137 is likely to represent the nucleophile (W_nuc_) that hydrolyzes ATP in F-actin. The O – Be – W_nuc_ angle in the structure is 144° (Fig. 4a), whereas an angle of >150° is required for efficient nucleophilic attack^29^. Inspection of the reconstruction revealed that the density for W_nuc_ is extended (Fig. 4e), indicating that the position of W_nuc_ is not fixed, allowing it to move into a position that brings the O – Be – W_nuc_ angle >150° while remaining hydrogen-bonded to Q137. In other words, W_nuc_ likely exchanges between hydrolysis competent and hydrolysis less competent configurations, perhaps contributing to the relatively low hydrolysis rate (0.3 s^-1^) ^5^ of F-actin.

Although Q137 positions W_nuc_ in close proximity to P*γ*, the Q137 side chain cannot accept a proton to act as a catalytic base for the hydrolysis. In the structure, we found no other amino acids that are close enough to W_nuc_ for a direct interaction. Instead, W_nuc_ resides at ∼4.2 Å from a neighboring water (W_bridge_) (Fig. 4a), which is not close enough to form a hydrogen bond, but the movement of W_nuc_ into a hydrolysis competent position would also place W_nuc_ in hydrogen bonding distance to W_bridge_. By forming hydrogen bonds with D154 and H161, W_bridge_ represents a Lewis base with a high potential to activate W_nuc_ and act as an initial proton acceptor during hydrolysis, followed by transfer of the proton to D154, as previously predicted by simulations^57, 58^, or alternatively, to H161. We conclude that Q137 coordinates W_nuc_ but that the hydrogen bond network comprising W_bridge_, D154 and H161 is responsible for the activation of W_nuc_ and proton transfer. Indeed, the ATP hydrolysis rates of the Q137 to alanine (Q137A) actin mutant are slower but not abolished^59^, whereas the triple mutant Q137A/D154A/H161A-actin exhibits no measurable ATPase activity^12^.

### Positions of water molecules determine polymerization and ATP hydrolysis kinetics

We next inspected the G- to F-actin transition in Ca^2+^-actin structures. The Ca^2+^-ion in G-actin is typically coordinated by the P*_β_* and P*γ* of ATP and five water molecules in a hepta-coordinated, pentagonal bipyramidal arrangement (Fig. 2d)^54, 60^. In contrast, in Ca^2+^-ADP-BeF_3_^-^ F-actin, the Ca^2+^ ion loses the interaction with one coordinating water and instead displays an octahedral coordination sphere (Fig. 2e). How does the G- to F-actin transition lead to changes in Ca^2+^-coordination? Globally, the flattening of Ca^2+^-actin triggers rearrangements that are analogous to those observed in Mg^2+^-actin (Fig. 3a, b), with a similar relocation of ordered water molecules from the narrowing SD3/1 cavity to the widening SD1 cavity (Fig. 3d, Movie S3). However, in Ca^2+^-G-actin, one of the relocating waters (W_x_) resides within the coordination sphere of Ca^2+^, indicating that the hydration shell of the Ca^2+^ ion needs to be altered for the G- to F-actin transition to occur. Thus, our analysis rationalizes why the inner-sphere coordination of Ca^2+^ changes from hepta-coordinated, pentagonal-bipyramidal in ATP-G-actin to hexa-coordinated, octahedral in ADP-BeF_3_^-^ F-actin (Fig. 1e, Fig. 2d, e Fig. 3d). Importantly, the required rearrangement of the Ca^2+^-coordination sphere could pose a kinetic barrier for the G- to F-actin transition, which provides a structural basis for the slower polymerization kinetics of Ca^2+^-actin compared to Mg^2+^-actin.

We also assessed the ATP hydrolysis mechanism of Ca^2+^-actin, which exhibits a 5x slower ATP hydrolysis rate (0.06 s^-1^) than Mg^2+^-actin (0.3 s^-1^)^5^. A structural comparison between Ca^2+^-ATP G-actin (pdb 1qz5) and Ca^2+^-ADP-BeF_3_^-^ F-actin demonstrates that the water corresponding to W_nuc_, which is hydrogen-bonded to Q137, locates closer to Be in F-actin (3.7 Å) than to P*γ* in G-actin (4.5 Å) (Fig. 4c, Supplementary Fig. 10c, d). Thus, the induction of ATP hydrolysis by a conformational change during polymerization is comparable in Ca^2+^-actin and Mg^2+^-actin. However, in Ca^2+^-ADP-BeF_3_^-^ F-actin, the distance between Q137 and W_nuc_ is 3.4 Å (3.2 Å in Mg^2+^-actin), and the O – Be – W_nuc_ angle is 137° (144° in Mg^2+^-actin) (Fig. 4e-g), making the position of W_nuc_ similar, but slightly less favorable for nucleophilic attack, providing a likely explanation for the slower hydrolysis rate of Ca^2+^-F-actin compared to Mg^2+^-F-actin.

**Fig. 4.**
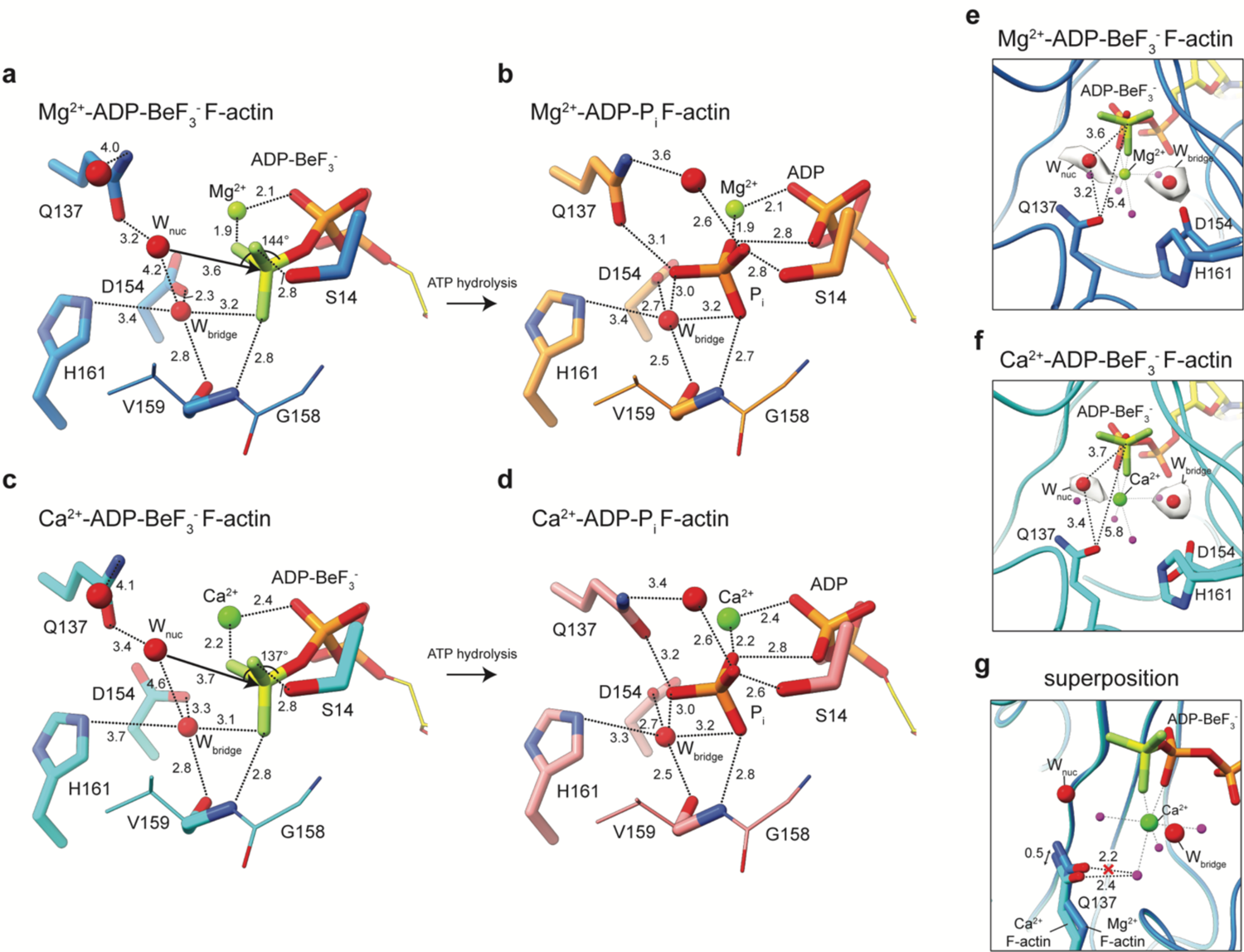
Mechanism of ATP hydrolysis. **a-d** Isolated amino acid and water arrangement near the nucleotide in Mg^2+^-ADP-BeF_3_^-^ F-actin **(a)**, Mg^2+^-ADP-P_i_ F-actin **(b)**, Ca^2+^-ADP-BeF_3_^-^ F-actin **(c)** and Ca^2+^-ADP-P_i_ F-actin **(d)**. All distances are shown in Å. Regions unimportant for interactions are depicted as smaller sticks. Amino acids and the proposed nucleophilic water (W_nuc_) and assisting water (W_bridge_) are annotated. **e,f** front view of the Pγ-mimic BeF_3_^-^ with densities for the putative W_nuc_ and W_bridge_ in structures of Mg^2+^-**(e)** and Ca^2+^-F-actin **(f)**. **g** Overlay of the amino-acid arrangement in front of ADP-BeF_3_^-^ in Mg^2+^-(blue) and Ca^2+^-F-actin (cyan). The nucleotide arrangement of Ca^2+^-F-actin is shown to emphasize that Q137 in its Mg^2+^-F-actin conformation would clash with a water in the inner-coordination sphere of the Ca^2+^-ion. The waters that coordinate the nucleotide-associated cation are colored magenta, whereas the water molecules important for the hydrolysis mechanism are shown as larger red spheres.

### P_i_ release from the F-actin interior occurs in a transient state

We next analyzed how ATP hydrolysis affects the F-actin nucleotide arrangement. In the Mg^2+^-ADP-P_i_ state, the cleaved P_i_ moiety is separated from ADP by at least 2.9 Å (Fig. 1c, Fig. 4b, Supplementary Fig. 9a), indicating that ADP and P_i_ do not form a covalent bond, as expected. The coordination shell of the Mg^2+^-ion is octahedral, resembling the arrangement in Mg^2+^-ADP-BeF_3_^-^ F-actin. In the Mg^2+^-ADP F-actin structure, which represents the filament state after P_i_ release, the P_i_ binding site is occupied by a water molecule (Fig. 1d, Supplementary Fig. 9a). Despite this rearrangement, the position of the Mg^2+^-ion does not change and its coordination remains octahedral. Because this observation also holds true for structures of monomeric Mg^2+^-ADP-G-actin^44, 61^ (Supplementary Fig. 9a), our structures show that the Mg^2+^-ion bound in the active site of actin resides at a fixed position beneath the P*_β_* moiety of the nucleotide.

After ATP hydrolysis in Ca^2+^-F-actin, the coordination of Ca^2+^ is also octahedral in the ADP-P_i_ state (Fig. 1f, Fig. 4d, Supplementary Fig. 9b). Interestingly, one coordinating water is replaced by the C=O side chain of Q137 (Supplementary Fig. 9b, 11). Finally, following P_i_ release, the Ca^2+^-ADP F-actin structure reveals that the Ca^2+^ ion surprisingly changes position so that it is directly coordinated by both the P*_α_* and P*_β_* of ADP (Fig. 1g, Supplementary Fig. 9b), and four water molecules in an octahedral arrangement. Thus, in contrast to Mg^2+^, the Ca^2+^ ion position is not fixed in F-actin and its coordination changes considerably during the ATPase cycle. This difference at the active site probably decreases the stability of the filament and could explain the higher depolymerization rates of Ca^2+^-actin compared to Mg^2+^-actin^18^.

P_i_ is thought to exit from the F-actin interior through the so called ‘back door’^62^, which is formed by the side chains of R177 and N111 and the backbones of methylated histidine 73 (H73) and G74 (Fig. 5, Supplementary Fig. 12). In this model, S14 switches rotameric position to change its hydrogen-bonding interaction from the backbone amide of G74 to the one G158, thereby allowing P_i_ to approach the back door, where R177 would mediate its exit^62^. Based on a lower-resolution reconstruction, the back door was proposed to be open in ADP F-actin^36^. However, S14 hydrogen-bonds with G74 and the back door is closed in our ∼2.2 Å structures of both the ADP-P_i_ and ADP states of Mg^2+^-F-actin (Fig. 5) and Ca^2+^-F-actin (Supplementary Fig. 12). Thus, unexpectedly, our structures reveal that the back door closes again after P_i_ release, indicating that the F-actin conformation that allows for the exit of P_i_ is a transient state. In fact, the proposed rotameric switch of S14 towards G158 alone would not result in an opened back door, which suggests that larger rearrangements are required for P_i_ release. Such a transient, high-energy F-actin state could be further explored by MD simulations or time-resolved cryo-EM^63^ in future research, guided by our structures as high-quality starting models.

**Fig. 5.**
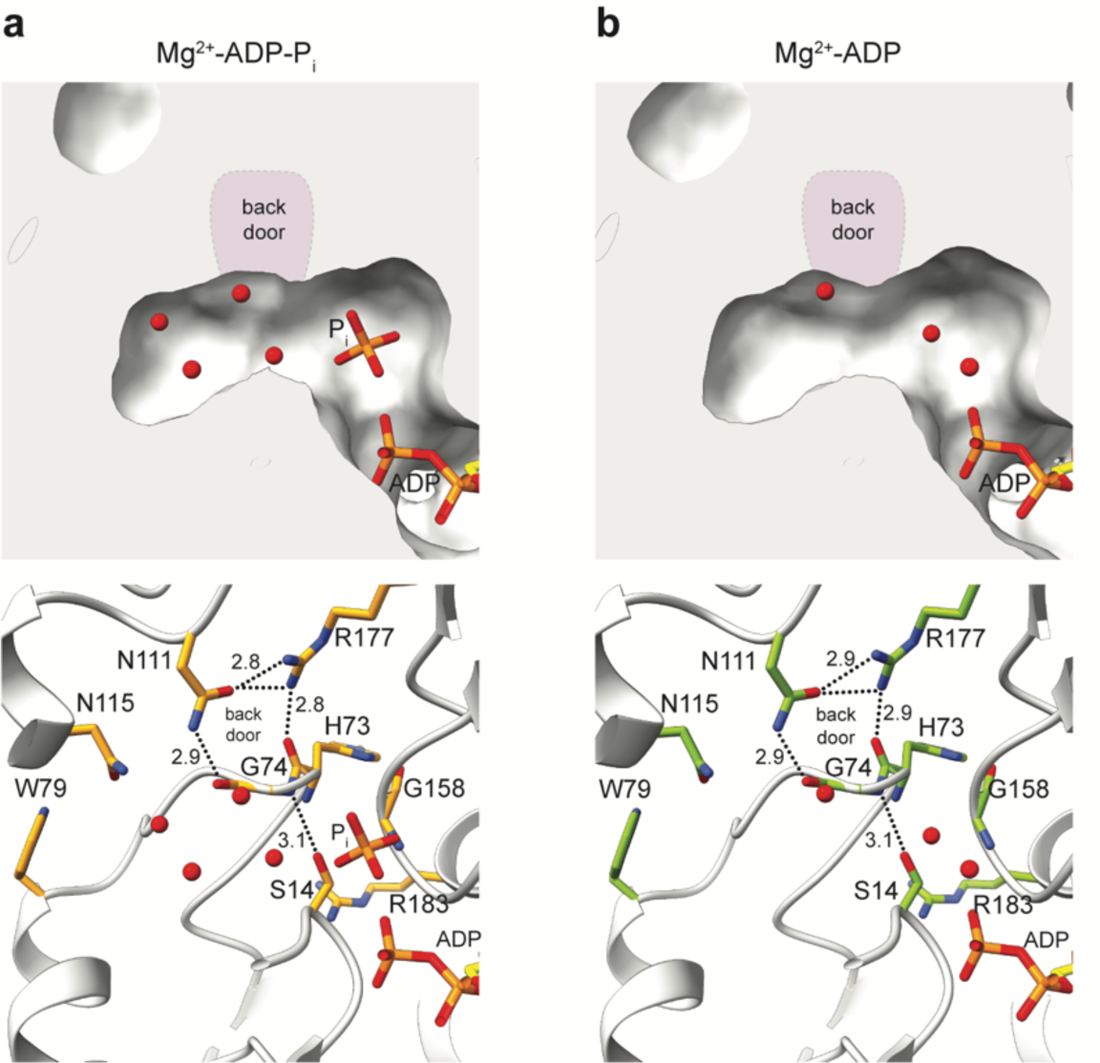
P_i_ release from the interior of Mg^2+^-F-actin. **a, b** Internal solvent cavities near the P_i_ binding site in ADP-P_i_ **(a)** and ADP **(b)** structures of Mg^2+^-F-actin. The upper panel shows the F-actin structure as surface with the bound P_i_ and water molecules. In the lower panel, F-actin is shown in cartoon representation, and the amino-acids forming the internal cavity are annotated and shown as sticks. The hydrogen bonds are depicted as dashed line. All distances are shown in Å. The position of the proposed back door is highlighted in purple in the upper panel. In none of the structures, the internal solvent cavity is connected to the exterior milieu.

### SD1 couples the nucleotide-binding site to the filament periphery

We next examined the global conformational changes that are associated with changes in the nucleotide states of F-actin. Biochemical and structural data have revealed the nucleotide-state dependent conformational mobility of the D-loop (residues 39–51) and the C-terminus at the intra-strand interface in the actin filament^35, 36, 43, 49^. The intra-strand arrangements in the current high-resolution Mg^2+^-F-actin structures are largely consistent with those in previous reconstructions, with a mixture of open/closed D-loop conformations in ‘young’ ATP-bound filaments and a predominantly closed D-loop arrangement in ‘aged’ ADP F-actin (Fig. 6). In the Mg^2+^-ADP-BeF_3_^-^ structure, we identified two intra-strand conformations that we could separate through a focused classification approach (Supplementary Fig. 13). The first conformation (∼37% of the particles, 2.38-Å resolution) represents the open D-loop, where the D-loop bends outwardly and interacts with the extended C-terminus of the adjacent actin subunit. In the second conformation (∼63% of the particles, 2.32-Å resolution), the C-terminus remains extended but turns away from the inwardly folded, closed D-loop (Fig. 6). In Mg^2+^-ADP-P_i_ F-actin, the C-terminus forms a compact, folded *α*-helix and the D-loop is predominantly closed. Finally, the Mg^2+^-ADP F-actin structure displays an extended C-terminus and a closed D-loop, thereby resembling the second conformation of the Mg^2+^-ADP-BeF_3_^-^ structure (Fig. 6). The intra-strand interfaces of the Ca^2+^ F-actin structures are very similar to those of the Mg^2+^-F-actin structures, but, surprisingly, the D-loop is predominantly in its closed state, even in the Ca^2+^-ADP-BeF_3_^-^ structure (Fig. 6).

**Fig. 6.**
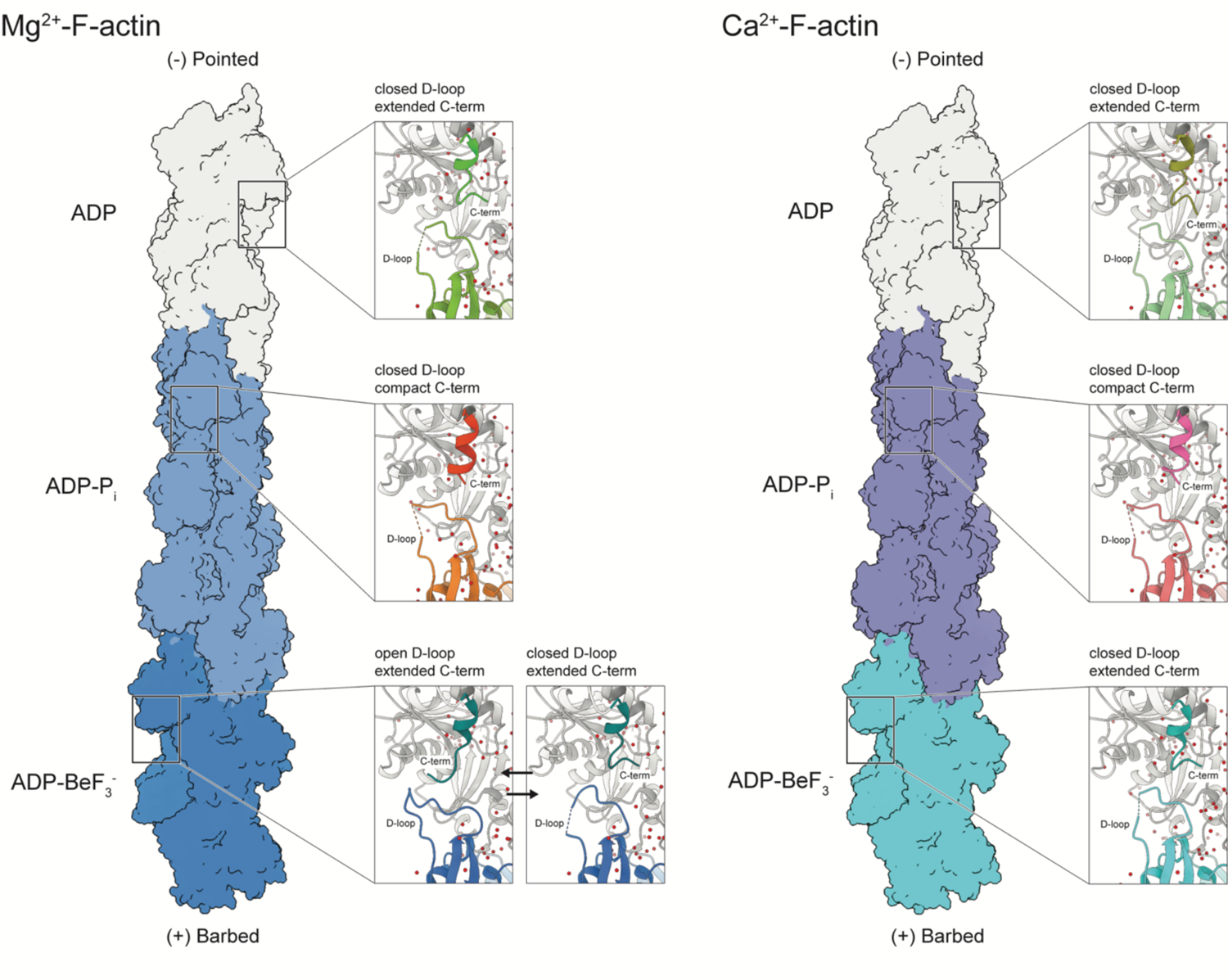
Arrangement of the F-actin intra-strand interface. Merged models of Mg^2+^-F-actin and Ca^2+^-F-actin in all nucleotide states. The structures are shown as surface and should be regarded as an infinitely long polymer. For each nucleotide state, a close-up of the observed intra-strand interface is depicted.

To understand how conformational changes in the nucleotide binding pocket are transmitted to the intra-strand interface at the filament surface, we analyzed the F-actin structures in detail. Surprisingly, we could not identify a direct communication path between the D-loop and the nucleotide binding site in our structures. In fact, a superposition of the Mg^2+^-ADP-BeF_3_^-^ structures with the separated open and closed D-loop conformations in the central actin subunit (Supplementary Fig. 13, 14), revealed that the only differences between the two structures, besides the D-loop, are not found in the central subunit, but instead in SD1 (including C-terminus) and SD3 of the adjacent actin subunit (Supplementary Fig. 14). Our data therefore suggest that the D-loop conformation in an actin subunit is not affected by changes in the same subunit, but rather by changes in SD1 of its neighbor subunit. Although our structures do not explain why the intra-strand interface can adopt two conformations in the ATP state of Mg^2+^-actin, we were able to identify the structural basis for the nucleotide-dependent conformation of the C-terminus. After ATP hydrolysis, Q137 in the nucleotide binding pocket moves upward by ∼0.4 Å in the ADP-P_i_ state so that it resides within 3.1 Å of P_i_ (Fig 4b, Fig. 7, Movie S4). This upward movement of Q137 triggers a sequence of small movements in the SD1; the proline-rich loop (residues 108 – 112) moves slightly forward and changes the position of E107, which forms a salt bridge with R116; the relocation of the E107-R116 salt bridge allows the penultimate residue C374 to flip into a hydrophobic pocket that is formed by Y133, I357 and V370; permitting R116 to interact with the carboxylate group of the C-terminal residue F375 (Fig. 7c, d, Movie S4). Altogether, these changes result in a compact, folded C-terminal helix, which then unfolds again when Q137 moves downward in the ADP state after P_i_ release (Supplementary Fig. 15). Since we also observed changes in the position of Q137 between the structures of Mg^2+^- and Ca^2+^-bound F-actin in the ADP-BeF_3_^-^ structures (Fig. 4g), these data suggest strongly that Q137 and its surrounding residues represent a major region within the interior of F-actin that is capable of sensing the nucleotide state and transmitting it to the periphery.

**Fig. 7.**
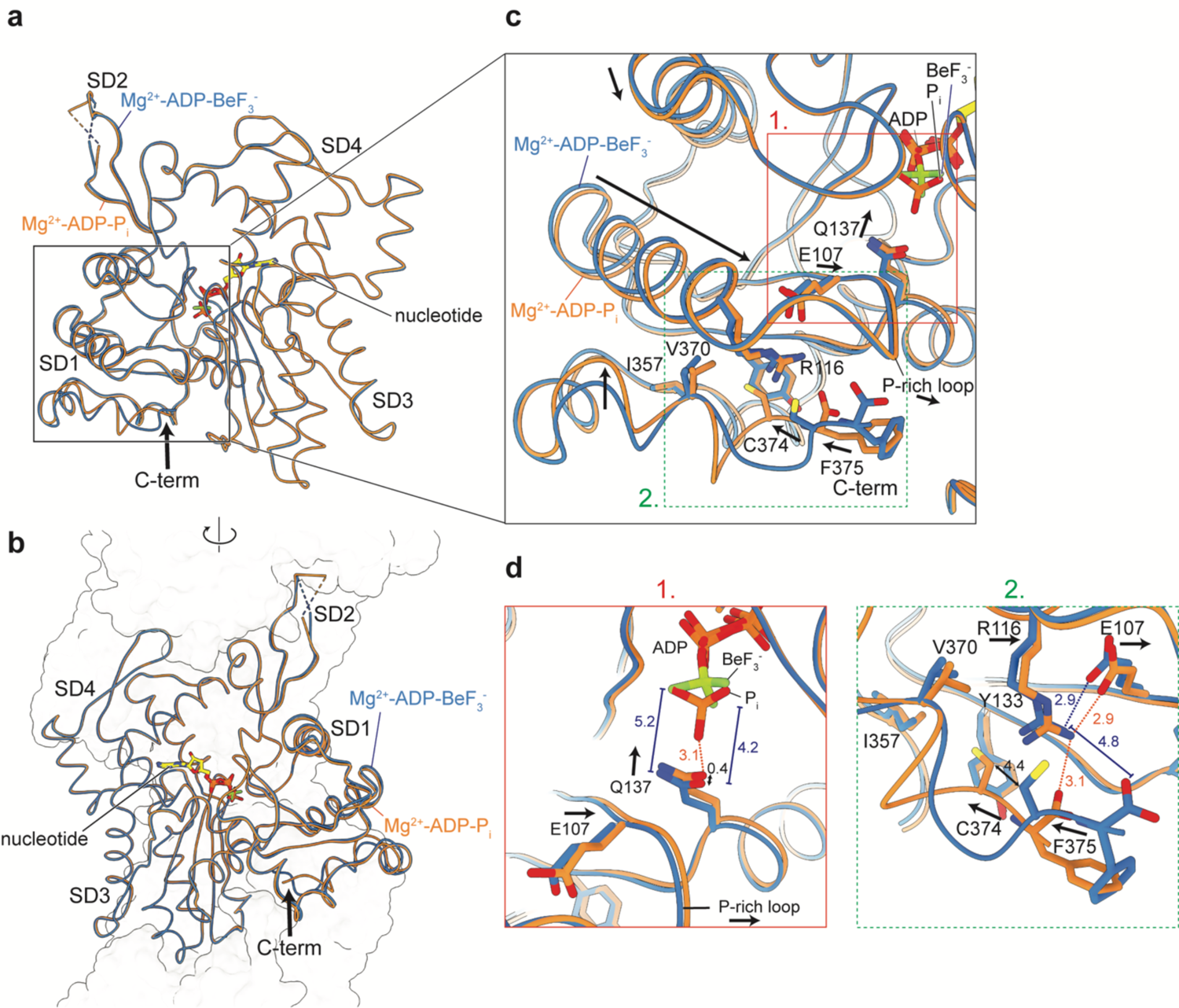
Structural coupling of the nucleotide binding site to the filament exterior. **a, b** Overlay of one actin subunit in the Mg^2+^-ADP-BeF_3_^-^ and Mg^2+^-ADP-P_i_ structures with annotated subdomains, shown in two orientations. The location of the C-terminus (C-term) is accentuated with a large arrow. For the Mg^2+^-ADP-BeF_3_^-^ structure, the closed D-loop conformation is shown as it represents the major intra-strand arrangement. In **(b)**, the surface-contour of other actin subunits within the filament is depicted. **c** Differences in the SD1 of F-actin in the Mg^2+^-ADP-BeF_3_^-^ and Mg^2+^-ADP-P_i_ structures. Residues thought to be important for the movement are annotated. **d** Zoom of the nucleotide binding site (1.) and C-terminal region (2.) of the SD1. In (**c)** and **(d)**, arrows depict the direction of the putative movement from the Mg^2+^-ADP-BeF_3_^-^ to the Mg-ADP-P_i_ structure. All distances are shown in Å. Distances shown in the Mg^2+^-ADP-BeF_3_^-^ structure are colored blue, whereas those in the Mg-ADP-P_i_ structure are colored orange.

### Cofilin senses the γ-phosphate moiety in F-actin

It has previously been proposed, by our group and others, that the intra-strand interface represents a major site for ABPs such as cofilin to sense the nucleotide state of F-actin. Cofilin binds and changes the helical twist of actin filaments by wedging itself between the C-terminus and D-loop^37, 38, 64, 65^, which, as we previously proposed, may be inhibited by the intra-strand conformation where the C-terminus interacts with the open D-loop^35^. In the structures of Ca^2+^-F-actin, we observed similar arrangements of the intra-strand interface compared to those of Mg^2+^-F-actin, except that the conformation where the C-terminus interacts with the open D-loop conformation is absent in Ca^2+^-ADP-BeF_3_^-^ F-actin (Fig. 6). We hypothesized that if the D-loop arrangement represents the dominant signal for cofilin to recognize the nucleotide state of the filament, cofilin would be capable of binding efficiently to Ca^2+^-F-actin regardless of the nucleotide state, since we only observed the closed D-loop in all Ca^2+^-F-actin structures (Fig. 6).

To assess this, we performed co-sedimentation assays of human cofilin-1 and F-actin, using the three nucleotide states of Ca^2+^-F-actin that we employed for the cryo-EM structure determination. Intriguingly, the assays revealed that human cofilin-1 only binds substantially to ADP-F-actin, and not to ADP-BeF_3_^-^ F-actin and ADP-P_i_ F-actin (Supplementary Fig. 16, 17). As the intra-strand interface in the Ca^2+^-ADP-BeF_3_^-^ F-actin and Ca^2+^-ADP F-actin structures are nearly identical (Fig. 6), the conformation of the C-terminus and D-loop cannot represent the only sensor for human cofilin-1 binding. What other mechanism does cofilin use to sense the nucleotide state? Our structures reveal that BeF_3_^-^ and P_i_ make hydrogen bonding interactions with S14 of SD1; and the backbones of G158 and V159 of the SD3 (Supplementary Fig. 18). Thus, in the ATP and ADP-P_i_ states of F-actin, the *γ*-phosphate moiety forms a bridge between the two subdomains, which is absent in the ADP state. Previous studies indicated that the tight binding of cofilin to F-actin necessitates a change in helical twist of the filament^38^, which involves the rotation of SD1 and 2 towards a position that is more similar to the conformation of G-actin^65^ (Supplementary Fig. 18a). This SD1 rotation involves the movement of the loop of S14, which is not possible when S14 is hydrogen bonded to the *γ*-phosphate of the nucleotide in ATP or ADP-P_i_ F-actin. Our results therefore support the previously proposed model that cofilin cannot form a strong complex with F-actin when the *γ*-phosphate moiety is present^37^. This is in agreement with numerous biochemical observations that members of the ADP/cofilin family can only sever actin filaments when P_i_ or BeF_3_^-^ is removed from the active site^15, 16, 66, 67^.

### Conclusions and perspectives

The structures of F-actin in the ATP, ADP-P_i_ and ADP states at ∼2.2-Å resolution reveal the filament architecture and the arrangement of the nucleotide-binding pocket in unmatched detail, allowing us to revise certain statements about the flexibility and stability of F-actin that have persisted in the literature. Traditionally, the structure of F-actin has been described as polymorphic with a variable helical twist^68, 69^, and the ‘aged’ ADP state of F-actin is regarded as a structurally destabilized form of the filament^70, 71^. In contrast, our current structural investigation defines that the bulk filament is remarkably similar in all solved nucleotide states, including the ADP state, and therefore strongly argue against a destabilized state that is characterized by large amino acid rearrangements. Accordingly, ADP-bound actin filaments do not sever spontaneously in the absence of cofilin. We therefore propose a change of terminology: ADP F-actin should not be regarded as a destabilized state, but rather as a ‘primed state’, which exhibits faster depolymerization rates at the filament ends and is sensitive to cofilin binding and severing due to the absence of the *γ*-phosphate moiety. Our high-resolution structures furthermore reveal the mechanism of ATP hydrolysis in F-actin and provide a basis for understanding why Ca^2+^-actin displays slower polymerization rates than Mg^2+^-actin. These molecular mechanisms depend on the positions of waters within the F-actin structure, emphasizing that high-resolution structures of the filament are crucial for explaining these important aspects of F-actin assembly and aging. Importantly, our optimized single-particle cryo-EM workflow now also paves the way for the structure determination of F-actin bound to ABPs at comparably high resolutions, which will enhance our understanding of how ABPs remodel the actin cytoskeleton. Finally, because F-actin adopts a central role in eukaryotic physiology, it represents a promising target for the development of small-molecule imaging probes^72^ and therapeutic agents^73^. The rational design of protein-binding small molecules is greatly facilitated by the availability of high-resolution structures of the target protein, because visible water molecules may aid in drug lead design^74^. We therefore envision that our solvent-molecule visualizing structures of F-actin will serve as high-quality templates for the development of actin-binding drugs and small molecules, which may be tailored for imaging and, perhaps, even therapeutic applications^75, 76^.

## Methods

### Protein purification

Skeletal α-actin was purified from rabbit muscle acetone powder through an established protocol that was described previously^27, 35, 77^. 0.5 g of frozen muscle acetone powder was thawed, resuspended in 10 ml G-buffer (5 mM Tris pH 7.5, 0.2 mM CaCl_2_, 0.2 mM ATP, 0.5 mM TCEP, 0.1 mM NaN_3_) and stirred for 25 minutes at 4 °C. The suspension was then filtered and the pellet was again resuspended in 10 ml G-buffer and subjected to the same stirring procedure. After filtering, the 20 ml of filtered solution was ultracentrifuged at 100,000*g* for 30 minutes to remove any remaining debris. The supernatant was collected and actin was polymerized by the addition of 2 mM MgCl_2_ and 100 mM KCl (final concentrations) for 1 hour at room temperature. To remove ABPs bound to actin, solid KCl was added to the solution to bring the KCl concentration to 800 mM and the mixture was incubated for 1 hour at room temperature. Then, the actin filaments were pelleted by ultracentrifugation at 100,000*g* for 2 hours and resuspended in 5 ml G-buffer. Actin was depolymerized by dialysis in 1 L G-buffer for 2 days, with one buffer exchange per day. On the third day, the solution was at ultracentrifuged 100,000*g* for 30 minutes and G-actin was recovered. The 2-day procedure of actin polymerization, high-salt wash and depolymerization by dialysis in G-buffer was repeated once more to ensure removal of all impurities and ABPs. After depolymerization, purified G-actin was flash frozen in liquid nitrogen in 50 µl aliquots at a concentration of ∼28 µM and stored at −80 °C until further use. Human cofilin-1 was purified as described previously^78^.

### Reconstitution of F-actin in different functional states

Structural studies were performed on rabbit skeletal α-actin, which is identical to human skeletal α-actin in amino-acid sequence. G-actin aliquots were thawed and ultracentrifuged for 1 hour at 100,000*g* to remove aggregates. For structures determined with Mg^2+^ as nucleotide-associated cation, G-actin (∼28 µM) was mixed with 0.5 mM EGTA and 0.2 mM MgCl_2_ to exchange Ca^2+^ for Mg^2+^ 5 – 10 minutes before polymerization. In all subsequent steps, buffers contained CaCl_2_ for the isolation of F-actin with Ca^2+^ as divalent cation, or MgCl_2_ for the isolation of F-actin with Mg^2+^ as divalent cation. Actin polymerization was induced by the addition of 100 mM KCl and 2 mM CaCl_2_/MgCl_2_ (final concentrations). Actin was polymerized at room temperature for 2 hours and subsequently overnight at 4 °C. The next morning, filaments were isolated through ultracentrifugation at 100,000*g* for 2 hours.

For the aged ADP F-actin states, the filament pellet was resuspended in F^-^ buffer: 5 mM Tris pH 7.5, 100 mM KCl, 2 mM CaCl_2_/MgCl_2_, 2 mM NaN_3_, 1 mM DTT. F-actin was used for cryo-EM sample preparation ∼1 hour after pellet resuspension.

For the ADP-BeF_3_^-^ states of F-actin, the filaments were resuspended in F^-^ buffer supplemented with 0.75 mM BeF_2_ and 5 mM NaF. Because the on-rate of BeF_3_^-^ for ADP-F-actin is relatively slow^43^, the filaments were incubated in this buffer for >6 hours prior to cryo-EM sample preparation to ensure saturation with BeF_3_^-^.

To isolate F-actin in the ADP-P_i_ state, we resuspended the actin pellet in F^-^ phosphate-buffer: 5 mM Tris, 50 mM KCl, 2 mM CaCl_2_/MgCl_2_, 2 mM NaN_3_, 1 mM DTT, 50 mM potassium phosphate pH 7.5. To remove any potential precipitates of calcium phosphate and magnesium phosphate, we filtered the buffers directly before use. The filaments were incubated in F^-^ phosphate-buffer for >6 hours prior to cryo-EM sample preparation.

### Cryo-EM grid preparation

2.8 µl of F-actin sample (3 – 16 µM) was pipetted onto a glow-discharged R2/1 Cu 300 mesh holey-carbon grid (Quantifoil). After incubating for 1 – 2 seconds, excess solution was blotted away and the grids were plunge frozen in liquid ethane or a liquid ethane/propane mixture using a Vitrobot Mark IV (Thermofisher Scientific). The Vitrobot was operated at 13 °C and the samples were blotted for 9 seconds with a blot force of −25.

### Cryo-EM grid screening and data collection

Grids were pre-screened on a 200 kV Talos Arctica Microscope (Thermofisher Scientific) equipped with a Falcon III detector (Thermofisher Scientific). Typically, low-magnification grid overviews (atlases) were collected using EPU (Thermofisher Scientific). Afterwards, ∼2 holes per grid square were imaged at high magnification for a total of 5 grid squares to visualize F-actin. The grids that displayed optimal filament concentration and distribution were then retrieved from the microscope and stored in auto grid boxes (Thermofisher Scientific) in liquid nitrogen until further use for high-resolution data collection.

All datasets were collected on a 300 kV Titan Krios microscope (Thermofisher Scientific) equipped with a K3 detector (Gatan) and a post-column energy filter (slit width: 15 eV). Movies were obtained in super-resolution at a pixel size of 0.3475 Å, with no objective aperture inserted. All datasets were collected on the same microscope at the same magnification of 130,000x, to ensure that the resulting cryo-EM density maps could be compared directly without issues caused by pixel size discrepancies. Using EPU, we collected ∼10,000 movies per dataset in 60 – 80 frames at a total electron exposure of ∼72 – 90 e^-^/Å^2^. The defocus values set in EPU ranged from −0.7 to −2.0 µm. The data quality was monitored live during acquisition using TranSPHIRE^79^. If necessary, the microscope was realigned to ensure optimal imaging conditions. An overview of the collection settings used for each dataset can be found in Table 1 and 2.

### Cryo-EM image processing

For each dataset, movies preprocessing was performed on the fly in TranSPHIRE^79^; the super-resolution movies were binned 2x (resulting pixel size: 0.695 Å), gain corrected and motion corrected using UCSF MotionCor2^80^, CTF estimations were performed with CTFFIND4.13^81^ and F-actin segments were picked using the filament picking procedure in SPHIRE_crYOLO^82, 83^ using a box distance of 40 pixels / 27.8 Å and a minimum number of six boxes per filament. The resulting particles were extracted in a 384×384 pixel box and further processed into the pipeline of helical SPHIREv1.4^84^. The number of extracted particles differed per dataset and ranged from 1,296,776 (Mg^2+^-ADP dataset) to 3,031,270 (Ca^2+^-ADP-P_i_ dataset) particles. For each dataset, the particles were 2D classified in 20k batches using ISAC2^85^ (sp_isac2.py). All classes were then pulled together and manually inspected and those that represented non-filament picks and ice contaminations were discarded. A virtual substack was created of the remaining particles (sp_pipe.py isac_substack) and the particles were subjected to 3D helical refinement^79^ using meridien alpha. This refinement approach within SPHIRE imposes helical restraints tailored to the helical sample to facilitate the refinement process, but does not actively apply helical symmetry.

Hence, symmetrization artifacts during refinement are avoided. We refined the F-actin structures with a restrained tilt angle during exhaustive search (--theta_min 90 –theta_max 90 –howmany 10) and used a filament width of 140 pix (97.3 Å) and a helical rise of 27.5 Å to limit shifts larger than one subunit to prevent duplication of particles. For the first processed dataset, EMD-11787^75^ was lowpass filtered to 25 Å and supplied as initial model for the refinement. The first meridien alpha refinement of each dataset was performed without a mask; the resulting 3D density map of this refinement was then used to create a soft mask using sp_mask.py that covered ∼85% of the filament (326 pixels in the Z-direction). The global refinement was then repeated with the same settings but with the mask applied. These masked refinements yielded F-actin reconstructions at resolutions of 2.6 – 3.0 Å. The particles where then converted to be compatible with Relion^86^ using sp_sphire2relion.py. Within Relion 3.1.0, the particles were subjected to Bayesian polishing^87^ for improved estimation of particle trajectories; and to CTF refinements^88^ to estimate per-particle defocus values and to correct for beam tilt, higher-order aberrations and anisotropic magnification. We then performed 3D classification without image alignment (8 classes, 25 iterations, tau2fudge 4) to remove particles that did not contribute high-resolution information to the reconstruction. Typically, one or two high resolution classes containing the majority of particles were selected and the other low-resolution classes were discarded. Finally, after removal of duplicates, this set of particles was subjected to a masked refinement with solvent flattening FSCs and only local searches (initial sampling 0.9°) in Relion using the map (lowpass filtered to 4.0 Å), mask and particle orientations determined from SPHIRE. These refinements yielded cryo-EM density maps at resolutions of 2.15 – 2.24 Å according to the gold-standard FSC=0.143 criterion. The final maps were sharpened with a negative B-factor and corrected with the modulation transfer function of the K3 detector. Local resolution estimations were performed in Relion.

To separate the closed and open D-loop conformations in the Mg^2+^-ADP-BeF_3_^-^ reconstruction, the good 2,228,553 particles of this dataset were subjected to a focused classification without image alignment in Relion. We created a soft mask around an inter-F-actin contact comprising the D-loop of the central actin subunit and the C-terminus of the subunit directly above. Initial attempts to separate the D-loop conformations into two classes using a single density map as initial model were unsuccessful, because all particles would end up in a single class with a mixed closed/open D-loop population. We therefore classified the particles into two classes using two references; the jasplakinolide-bound, Mg^2+^-ADP-P_i_ (in-house structure) and Mg^2+^-ADP structures as templates for, respectively, open and closed D-loop conformations. After optimization of the tau2fudge parameter, this classification without image alignment (2 classes, 25 iterations, tau2fudge 500) yielded two classes with clearly distinguishable D-loop conformations. The particles belonging to each class (834,110 for the open D-loop, 1,394,443 for the closed D-loop) were then selected and separately refined using the map with mixed D-loop conformation (filtered to 4.0 Å) as reference and the SPHIRE-mask covering 85% of the filament. The resulting maps of the open D-loop (2.38 Å) and closed D-loop (2.32 Å) particles showed, respectively, the expected open and closed D-loop conformations at the region that was used for focused classification. The D-loop conformations remained a mix between open and closed in actin subunits within the map that were not used for focused classification.

### Model building, refinement and analysis

To build the F-actin models in the high-resolution density maps, the structure of F-actin in complex with an optojasp in the cis state^75^ (pdb 7AHN) was rigid-body fitted into the map of F-actin Ca^2+^-ADP state. We modelled five actin subunits in each map to capture the entire interaction interface within the filament, because the central subunit interacts with four neighboring protomers. The central actin subunit in the map was rebuilt manually in Coot^89^, and the other actin subunits were adjusted in Coot by applying non-crystallographic symmetry (NCS) using the central subunit as master chain. The structure was then iteratively refined using Coot (manually) and phenix real-space refine^90^ with NCS restraints but without imposing any geometry restraints. The structures of all other states were built by rigid-body fitting of the Ca^2+^-ADP structure in the map belonging to each F-actin state, followed by manual adjustments in Coot. These structures were then refined through a similar protocol of iterative cycles in Coot and phenix real-space refine. All solvent molecules (ions, waters) were placed manually in Coot in the central actin subunit, and were then placed in the other subunits using NCS. Because the local resolution of each F-actin reconstruction is highest in the center and lower at the periphery of the map, we inspected all waters in each structure manually before the final phenix refinement; water molecular with poor corresponding cryo-EM density were removed. A summary of the refinement quality for each structure is provided in Table 1 and 2. For the structural analysis, the central actin subunit in the structure was used, unless stated otherwise. The solvent cavities in the structures were calculated using the CASTp 3.0 web server^91^.

### Cofilin co-sedimentation assays

F-actin in different nucleotide states was prepared as for cryo-EM experiments (see above). Co-sedimentation assays were performed in 20-μl volumes by incubating 5 μM of F-actin with 5 μM of cofilin for 30 minutes at room temperature, and then by centrifuging the samples at 120,000 g in a TLA120.1 rotor for 15 minutes at 4°C. After centrifugation, aliquots of the supernatant and pellet fractions were separated by SDS-PAGE and analyzed by densitometry using Image Lab software, version 6.0.1 (Bio-Rad). The amount of cofilin bound to actin was calculated by subtracting the amount of cofilin that pelleted in the absence of F-actin from the amount of the ABP bound to F-actin. The data points are available in supplementary data. Statistical analysis was performed by ordinary one-way ANOVA test in Prism version 9 (GraphPad Software).

### Author contributions

S.R. conceived and supervised the study. B.U.K. and S.P. optimized data collection strategies. W.O. collected and processed all cryo-EM data and built the atomic models. W.O., S.P. and S.R. analyzed the structures. A.B. performed the cofilin co-sedimentation assays. W.O. and S.R wrote the manuscript, with critical input from all authors.

## Acknowledgements

We gratefully thank D. Prumbaum and O. Hofnagel for the assistance with cryo-EM data collection; S. Bergbrede for technical support in the wet lab; and T.D. Pollard, F. Merino and R.S. Goody for critical proofreading of the manuscript. We acknowledge W. Linke and A. Unger for supplying us with muscle acetone powder. This work was supported by funds from the Max Planck Society (to S.R.) and the European Research Council under the European Union’s Horizon 2020 Programme (ERC-2019-SyG, grant no. 856118 to S.R). A.B. was supported by an EMBO long-term fellowship. W.O. is a fellow of the Alexander von Humboldt foundation.

## Data availability

Data supporting the findings of this manuscript are available from the corresponding author (S.R.) upon reasonable request. The cryo-EM reconstructions will be deposited to the Electron Microscopy Data Bank before final publication, and the protein models will be uploaded to the Protein Data Bank.

---

**Supplementary Fig. 1.**
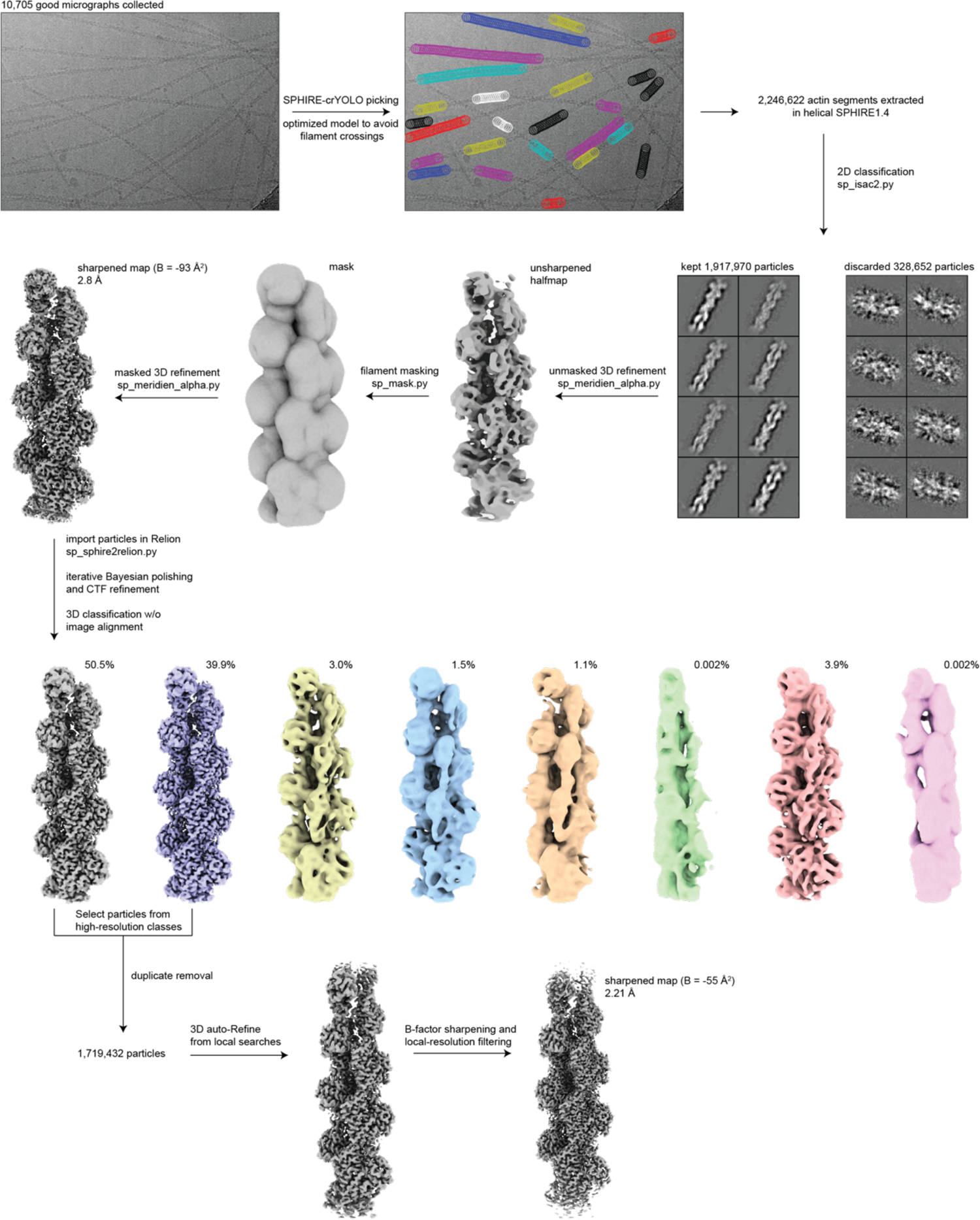
Cryo-EM image processing workflow. The image processing workflow that was used for all collected datasets is shown, with the Ca^2+^-ADP-BeF_3_^-^ F-actin dataset as example. All maps are shown in the same orientation.

**Supplementary Fig. 2.**
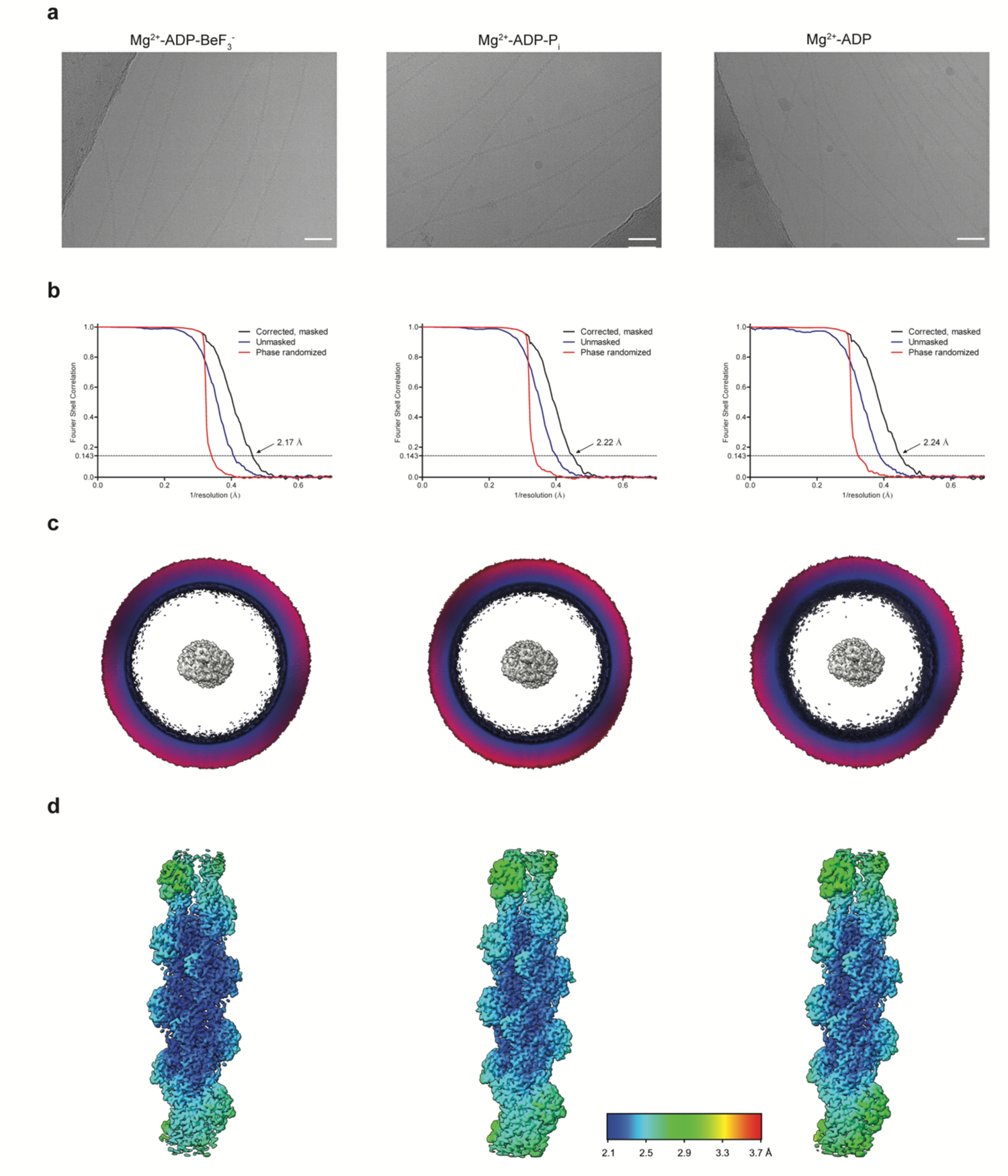
Processing of the Mg^2+^-F-actin cryo-EM datasets. **a** Micrograph depicting Mg^2+^-actin filaments in the ADP-BeF_3_^-^ (−2.1 μm), ADP-P_i_ (−2.4 μm) and ADP (−1.3 μm) states distributed in vitreous ice (defocus values between brackets). The scale bar is 400 Å. **b** Fourier-shell correlation plots for each Mg^2+^-F-actin structure of gold-standard refined masked (black), unmasked (blue) and high-resolution phase randomized (red) half maps. The FSC = 0.143 threshold is depicted as a dashed line. **c** Angular distribution of the particles used in the final reconstruction, shown along the filament axis. **d** Local-resolution estimations of the Mg^2+^-F-actin reconstructions, computed through Relion.

**Supplementary Fig. 3.**
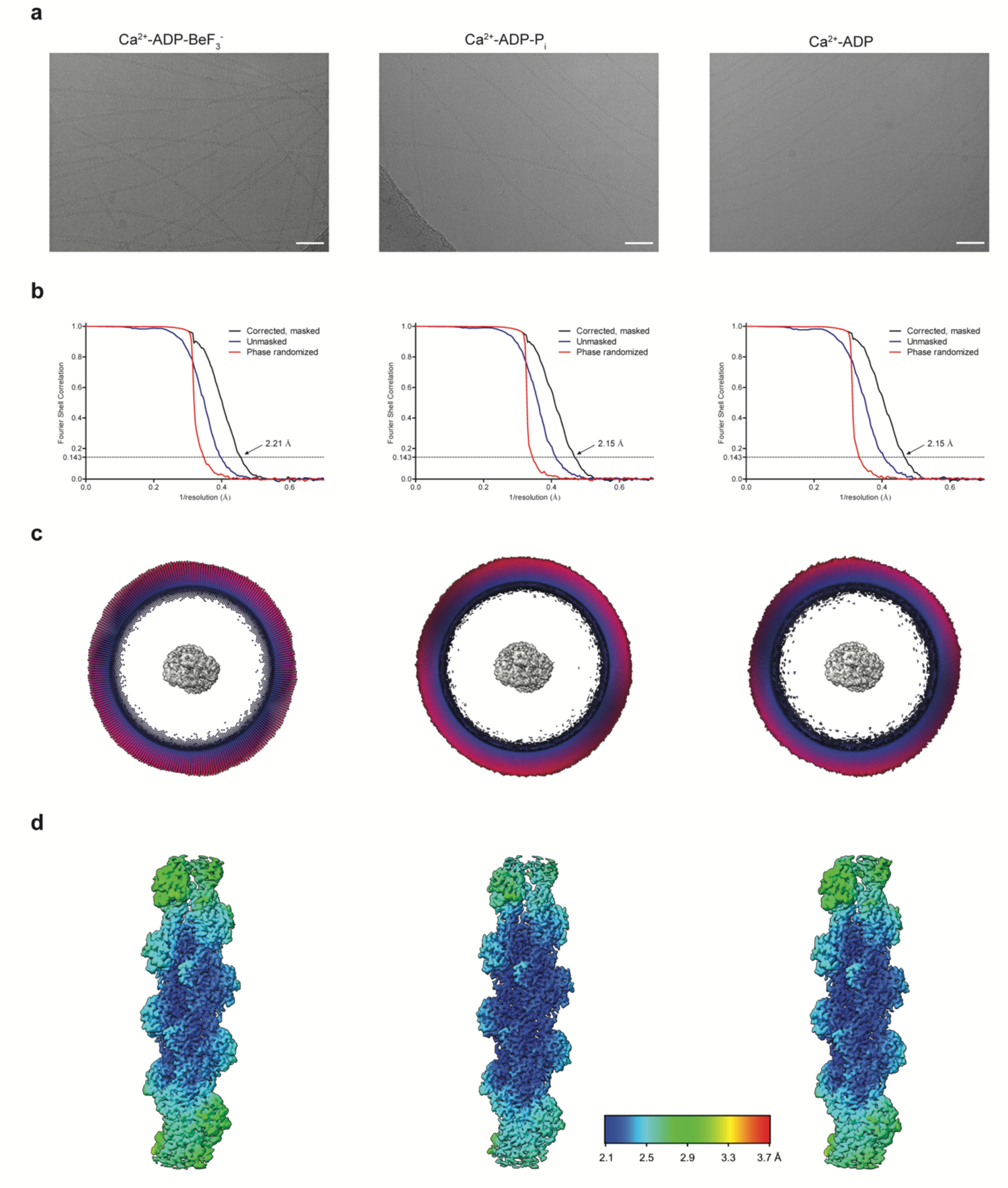
Processing of the Ca^2+^-F-actin cryo-EM datasets. **a** Micrograph depicting Ca^2+^-actin filaments in the ADP-BeF_3_^-^ (−1.4 μm), ADP-P_i_ (−2.0 μm) and ADP (−1.0 μm) states distributed in vitreous ice (defocus values between brackets). The scale bar is 400 Å. **b** Fourier-shell correlation plots for each Ca^2+^-F-actin reconstruction of gold-standard refined masked (black), unmasked (blue) and high-resolution phase randomized (red) half maps. The FSC = 0.143 threshold is depicted as a dashed line. **c** Angular distribution of the particles used in the final reconstruction, shown along the filament axis. **d** Local-resolution estimations of the Ca^2+^-F-actin reconstructions, computed through Relion.

**Supplementary Fig. 4.**
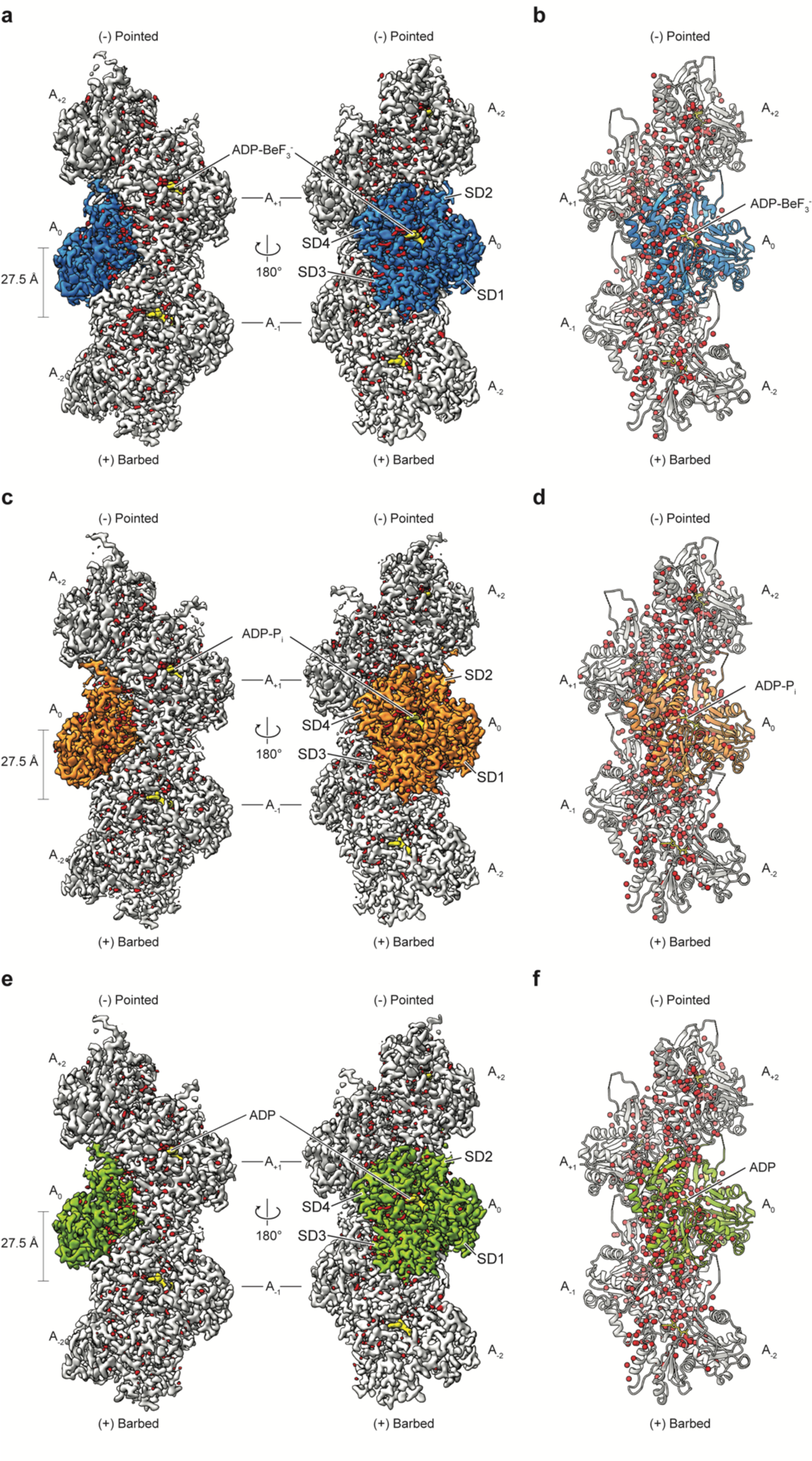
High resolution cryo-EM structures of Mg^2+^-F-actin allow for the modeling of water molecules. **a, c, e** Local-resolution filtered, sharpened cryo-EM density map of Mg^2+^-ADP-BeF_3_^-^ **(a)**, Mg^2+^-ADP-P_i_ **(c)** and Mg^2+^-ADP-F-actin **(e)** shown in two orientations. The subunits are labeled based on their location along the filament, ranging from the barbed (A_-2_) to the pointed (A_2_) end. The central actin subunit (A_0_) is colored blue **(a)**, orange **(c)** or green **(e)** the other four subunits are grey. Densities corresponding to water molecules are colored red. **b, d, f** Cartoon representation of the of Mg^2+^-ADP-BeF_3_^-^ **(b)**, Mg^2+^-ADP-P_i_-**(d)** and Mg^2+^-ADP F-actin **(f)** structures. Waters are shown as spheres to emphasize their location.

**Supplementary Fig. 5.**
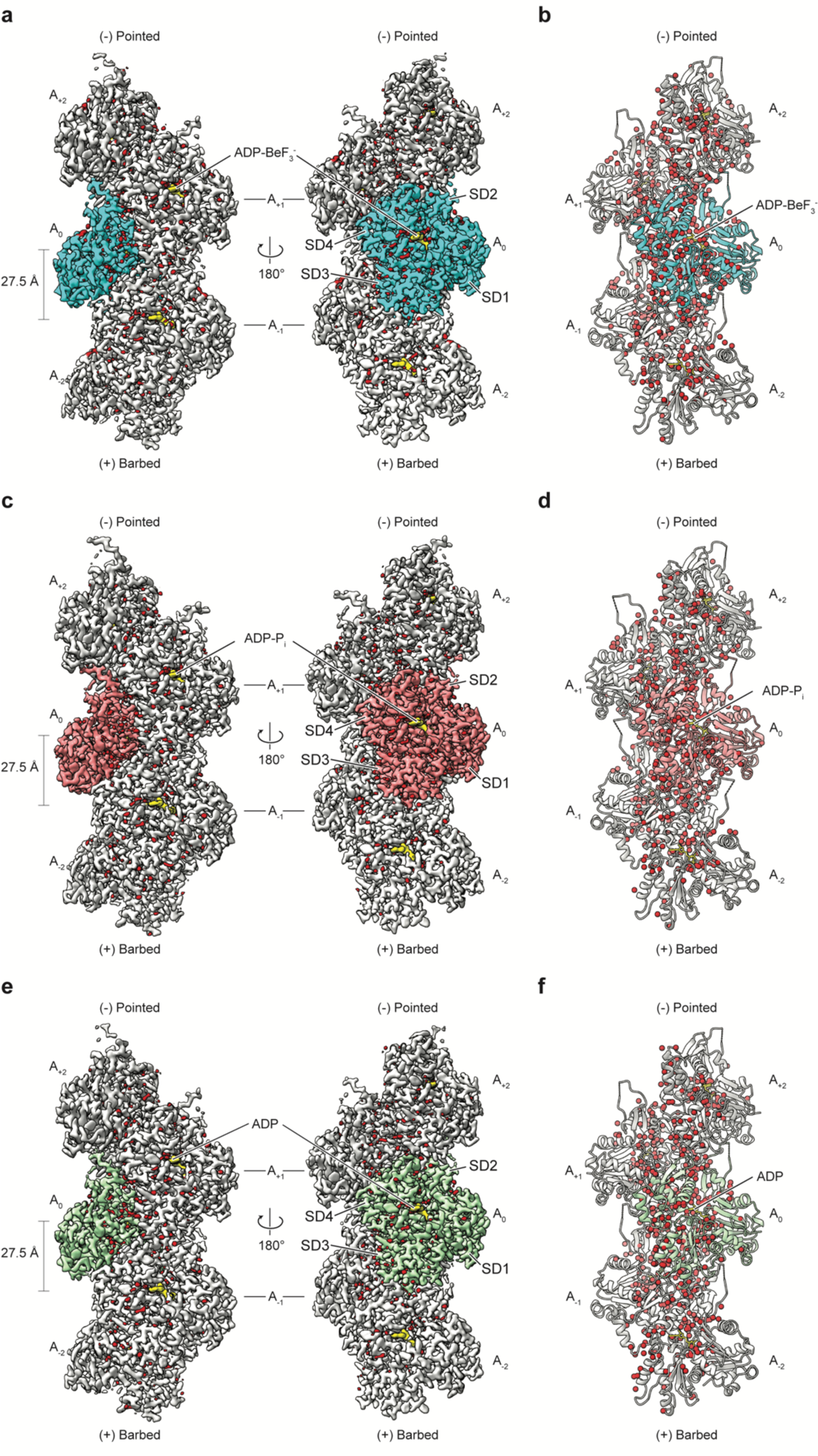
High-resolution cryo-EM structures of Ca^2+^-F-actin allow for the modeling of water molecules. **a, c, e** Local-resolution filtered, sharpened cryo-EM density map of Ca^2+^-ADP-BeF_3_^-^ **(a)**, Ca^2+^-ADP-P_i_ **(c)** and Ca^2+^-ADP F-actin **(e)** shown in two orientations. The subunits are labeled based on their location along the filament, ranging from the barbed (A_-2_) to the pointed (A_2_) end. The central actin subunit (A_0_) is colored cyan **(a)**, salmon **(c)** or palegreen **(e)** the other four subunits are grey. Densities corresponding to water molecules are colored red. **b, d, f** Cartoon representation of the of Ca^2+^-ADP-BeF_3_^-^ **(b)**, Ca^2+^-ADP-P_i_-**(d)** and Ca^2+^-ADP F-actin **(f)** structures. Waters are shown as spheres to emphasize their location.

**Supplementary Fig. 6.**
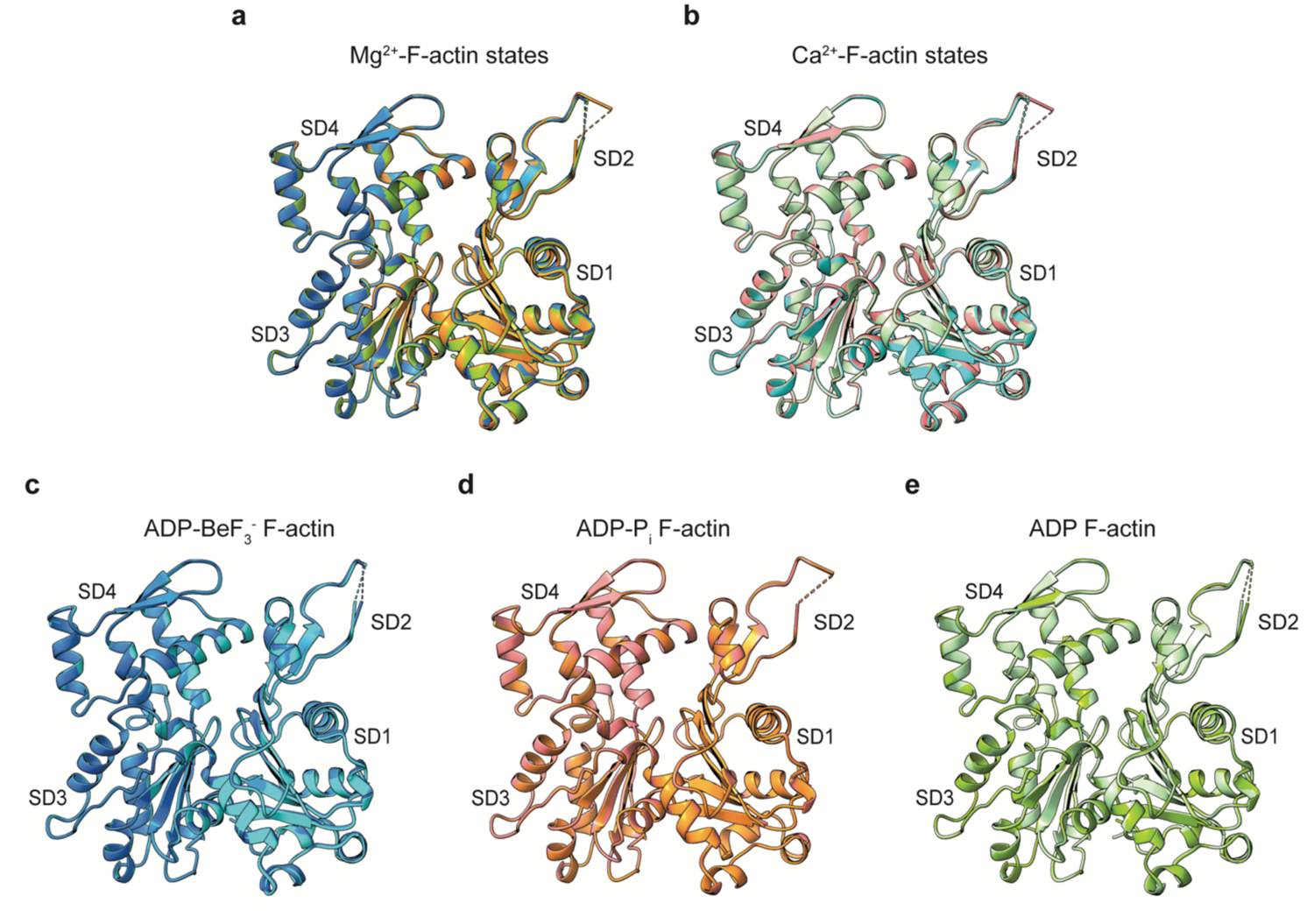
Structural similarities between high-resolution F-actin structures. **a** Superimposition of a single subunit of Mg^2+^-F-actin in ADP-BeF_3_^-^ (blue), ADP-P_i_ (orange) and ADP (green) states. **b** Superimposition of a single subunit of Ca^2+^-F-actin in ADP-BeF_3_^-^ (cyan), ADP-P_i_ (salmon) and ADP (pale-green) states. **c-e** Superimpositions of Mg^2+^-F-actin and Ca^2+^-F-actin in the ADP-BeF_3_^-^ **(c)**, ADP-P_i_ **(d)** and ADP **(e)** states. The coloring is consistent with the descriptions of **a** and **b**. For each overlay, the subdomains of actin (SD1 – SD4 are annotated).

**Supplementary Fig. 7.**
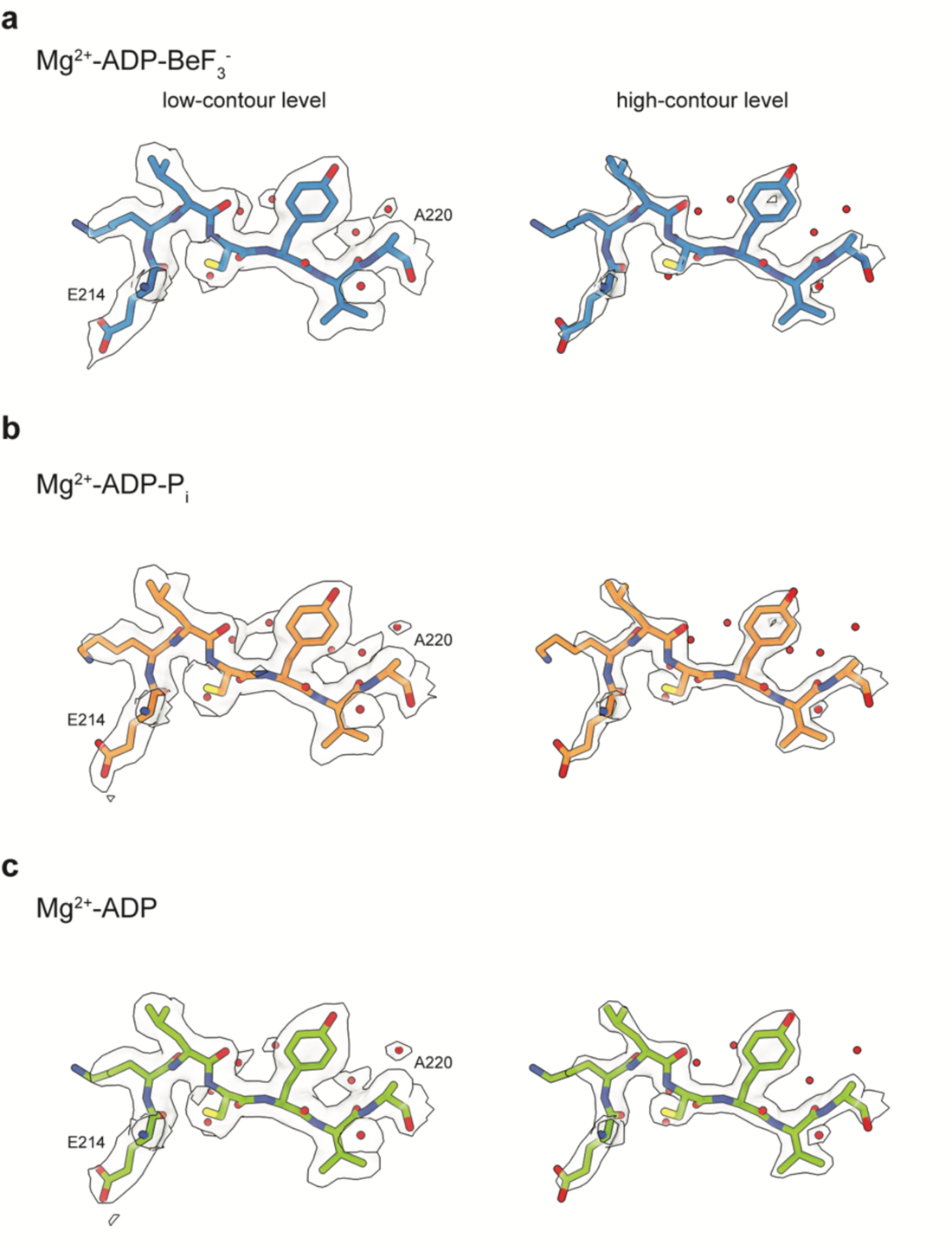
Modeling of selected regions in Mg^2+^-F-actin structures. **a-c** Cryo-EM density of residues E214 – A220 with modelled amino acids and water molecules of Mg^2+^-F-actin in the ADP-BeF_3_^-^ **(a)**, ADP-P_i_ **(b)** and ADP **(c)** states, shown at two different contour levels.

**Supplementary Fig. 8.**
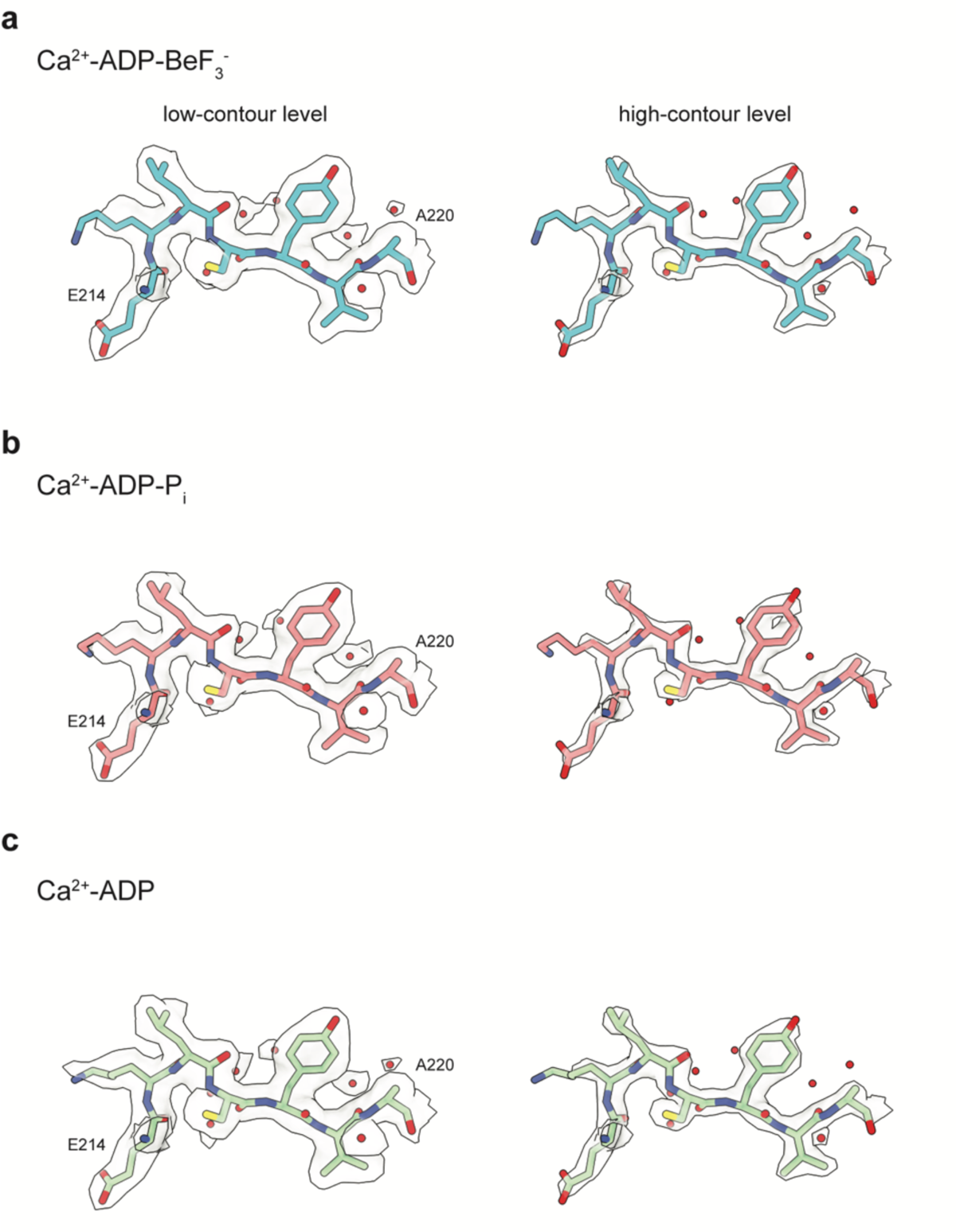
Modeling of selected regions in Ca^2+^-F-actin structures. **a-c** Cryo-EM density of residues E214 – A220 with modelled amino acids and water molecules of Ca^2+^-F-actin in the ADP-BeF_3_^-^ **(a)**, ADP-P_i_ **(b)** and ADP **(c)** states, shown at two different contour levels.

**Supplementary Fig. 9.**
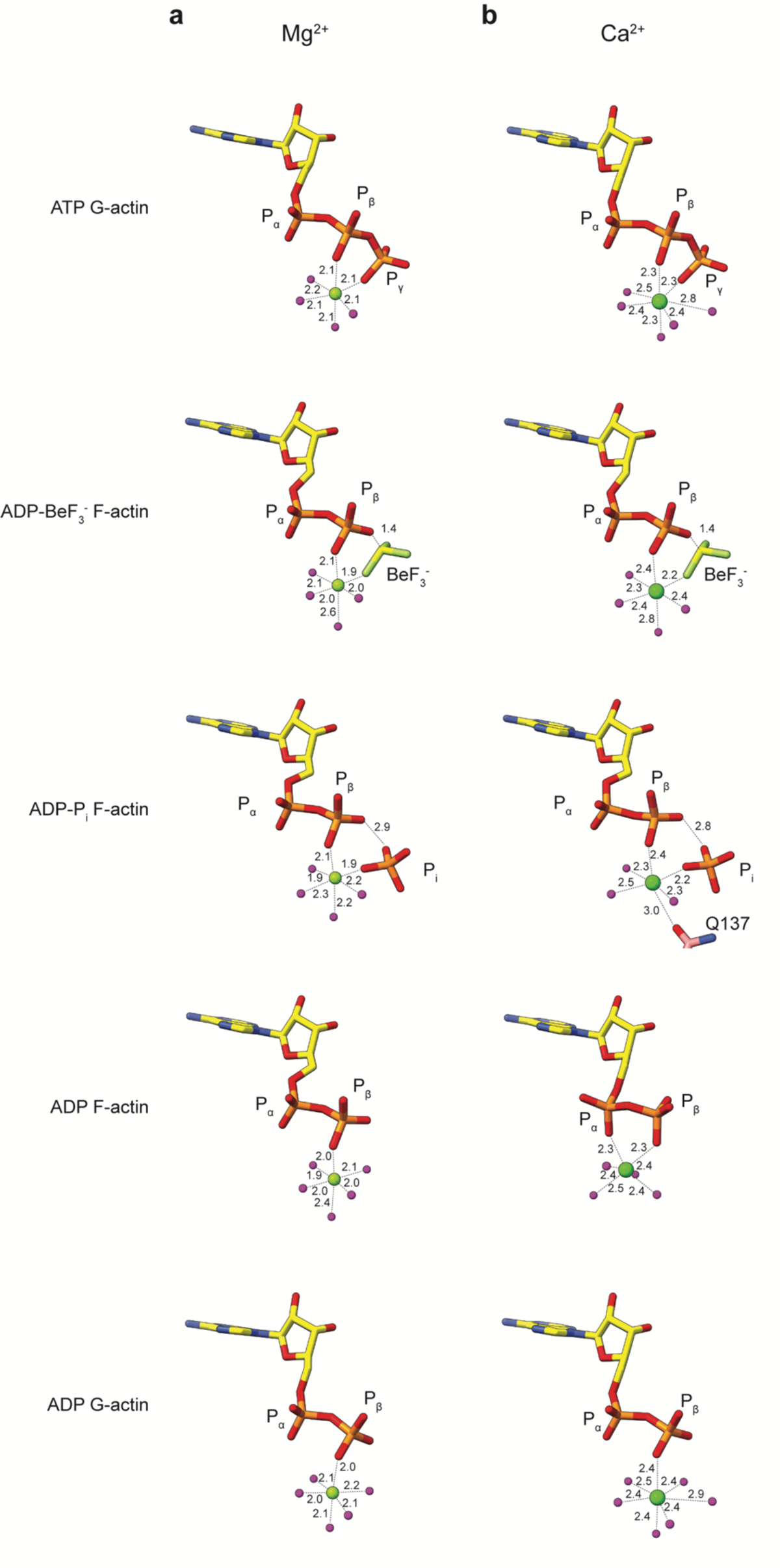
Ion coordination at the nucleotide-binding sites. **a, b** Nucleotide conformation and inner-coordination sphere of the cation for Mg^2+^-**(a)** and Ca^2+^-actin **(b)**. The shown G-actin models were selected from high-resolution crystal structures of rabbit G-actin in the following states: Mg^2+-^ATP (pdb 2v52, 1.45 Å), Mg^2+^-ADP (pdb 6rsw, 1.95 Å), Ca^2+^-ATP (pdb 1qz5, 1.45 Å) and Ca^2+^-ADP (pdb 1j6z, 1.54 Å). All distances are shown in Å. The distances between the cation and the molecules in its coordination sphere were not restrained during model refinement and may therefore deviate from ideal values.

**Supplementary Fig. 10.**
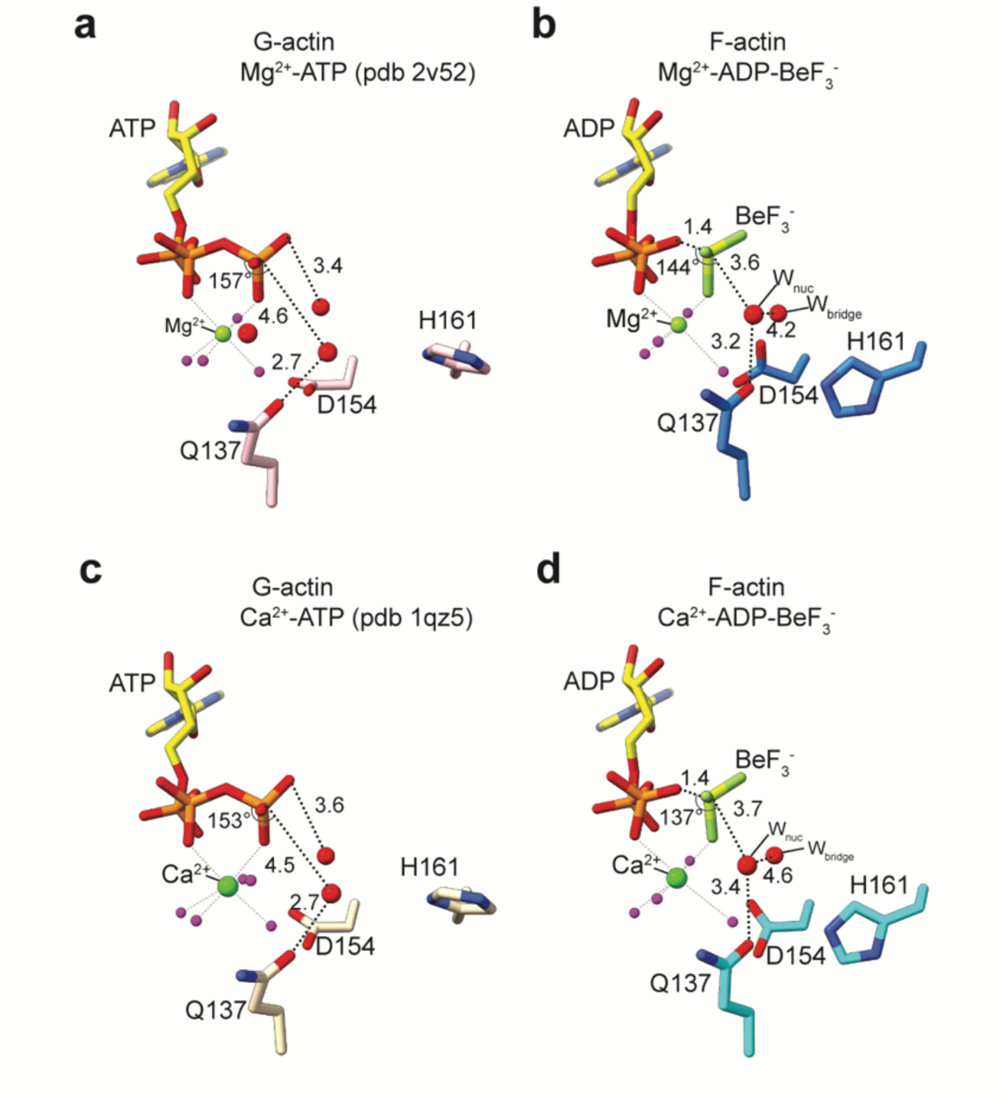
Increased ATPase activity of F-actin. **a-d** Water arrangement in front of the nucleotide in structures of Mg^2+^-ATP G-actin (pdb 2v52) **(a)**, Mg^2+^-ADP-BeF_3_^-^ F-actin **(b)**, Ca^2+^-ATP G-actin (pdb 1qz5) **(c)** and Ca^2+^-ADP-BeF_3_^-^ F-actin **(d)**. All distances are shown in Å. The waters that coordinate the nucleotide-associated cation are colored magenta, whereas the waters important for the hydrolysis mechanism are shown as larger red spheres.

**Supplementary Fig. 11.**
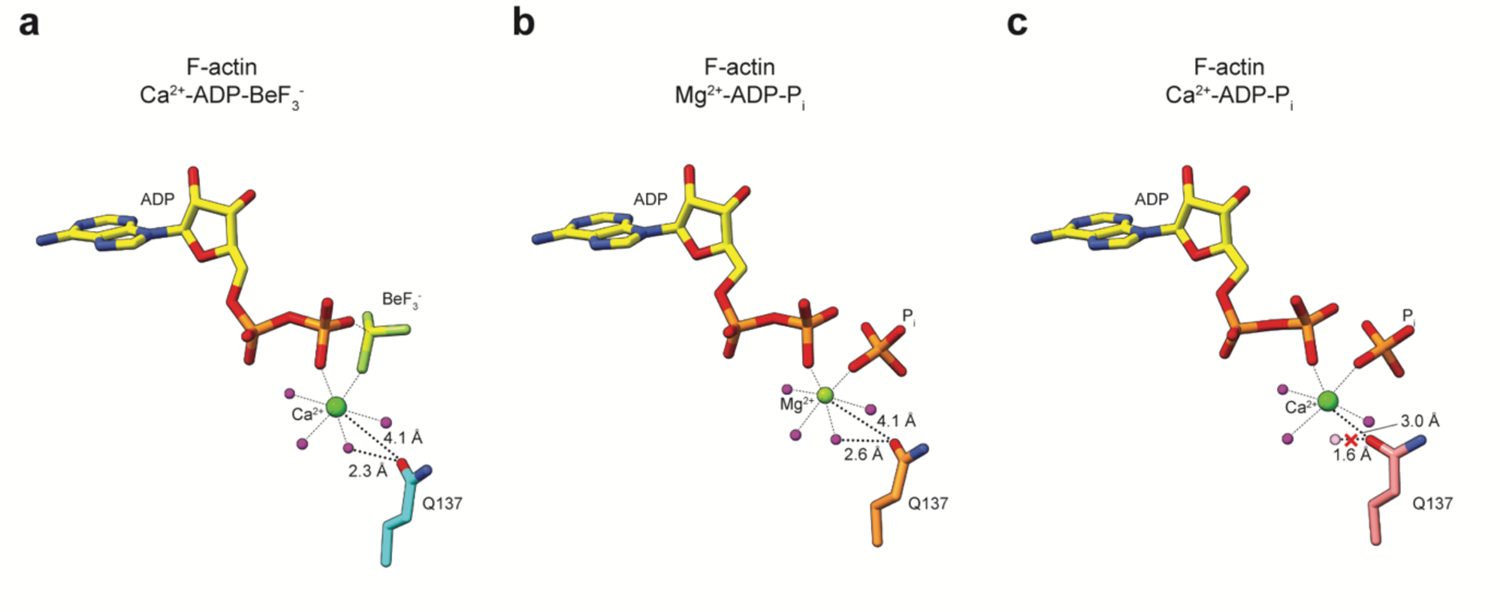
Change of Ca^2+^-coordination sphere in the Ca^2+^-ADP-P_i_ state. **a-c** Position of the nucleotide, cation and associated waters with respect to residue Q137 in the Ca^2+^-ADP-BeF_3_^-^ **(a)**, Mg^2+^-ADP-P_i_ **(b)** and Ca^2+^-ADP-P_i_ **(c)** states of F-actin. In the Ca^2+^-ADP-P_i_ state (panel **c**), the position of Q137 prevents the binding of one of the Ca^2+^-coordinating waters (shown in semi-transparent magenta), yielding an octahedral inner-coordination sphere of the Ca^2+^ ion with one missing water, but instead a coordination by Q137.

**Supplementary Fig. 12.**
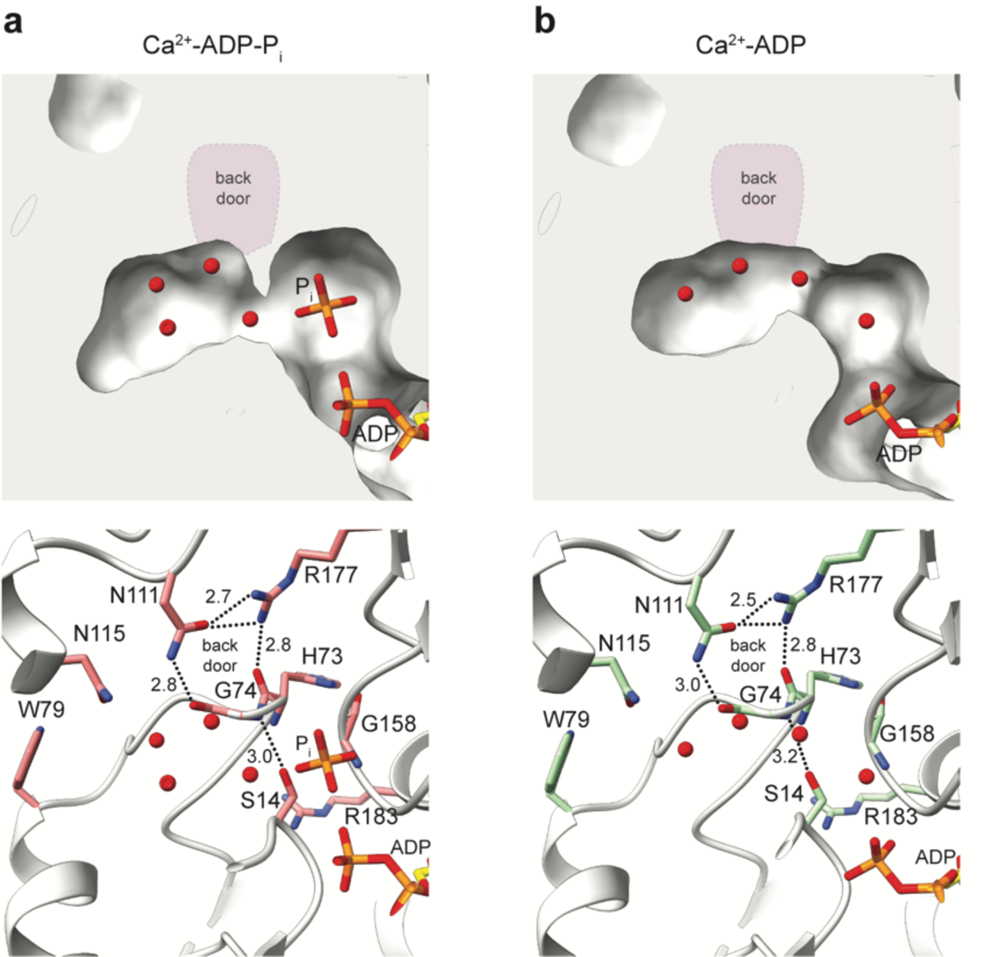
P_i_ release from the interior of Ca^2+^-F-actin. **a, b** Internal solvent cavities near the P_i_ binding site in ADP-P_i_ **(a)** and ADP **(b)** structures of Ca^2+^-F-actin. The upper panel shows the F-actin structure as surface with the bound P_i_ and water molecules. In the lower panel, F-actin is shown in cartoon representation, and the amino-acids forming the internal cavity are annotated and shown as sticks. The hydrogen bonds are depicted as dashed line. All distances are shown in Å. The position of the proposed back door is highlighted in purple in the upper panel. In none of the structures, the internal solvent cavity is connected to the exterior milieu.

**Supplementary Fig. 13.**
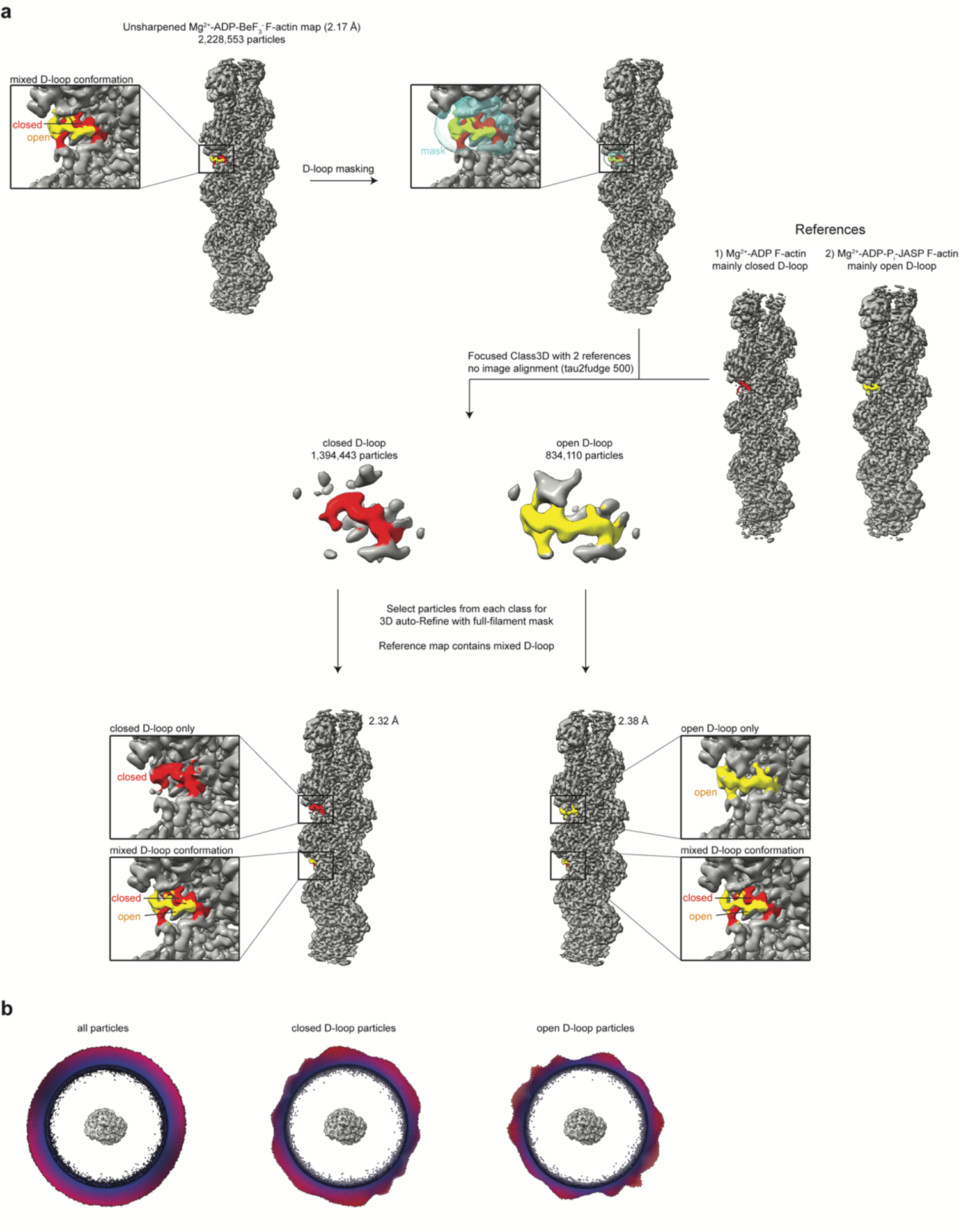
Focused classification of two D-loop conformations. **a** Focused classification strategy for the separation of the D-loop states of the Mg^2+^-ADP-BeF_3_^-^ F-actin dataset. The closed D-loop is colored red, whereas the open D-loop is colored yellow. **b** Angular distribution of all particles used in the reconstruction of the Mg^2+^-ADP-BeF_3_^-^ structure (left); and of the particles used for the reconstructions of the isolated closed (middle) and open (right) D-loop.

**Supplementary Fig. 14.**
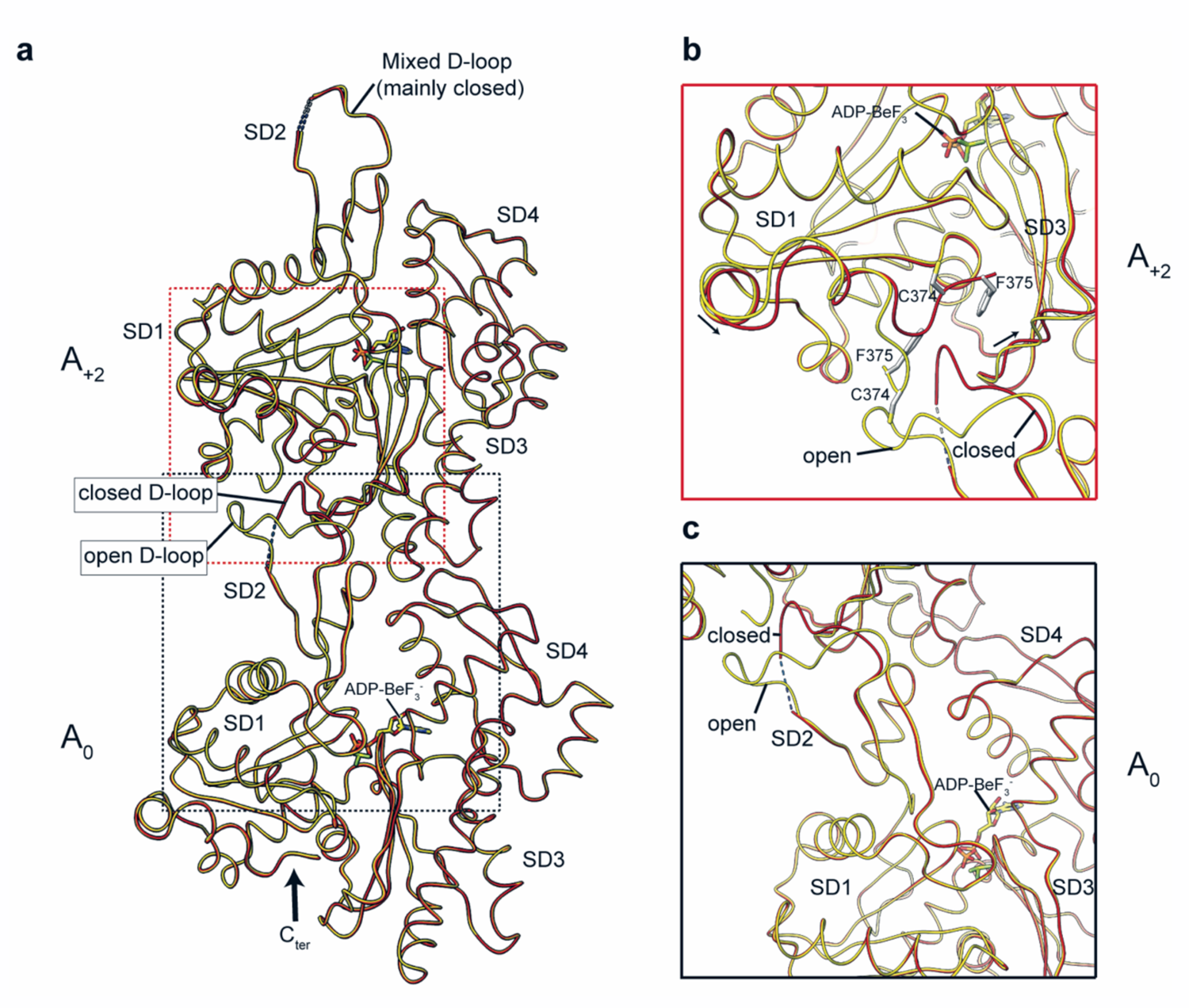
Conformational differences associated with open and closed D-loop. **a** superposition of the refined Mg^2+^-ADP-BeF_3_^-^ F-actin structures with separated D-loop conformations in the central actin subunit (A_0_). For both the open (yellow) and closed (red) D-loop states, the A_0_ and A_+2_ actin subunits are shown. **b, c** Close up of the A_0_ **(b)** and A_+2_ **(c)** subunits. In panel **b**, arrows depict the movements in the SD1 and SD3 of the A_+2_ subunit associated with the change of D-loop conformation from open to closed in the central A_0_ subunit.

**Supplementary Fig. 15.**
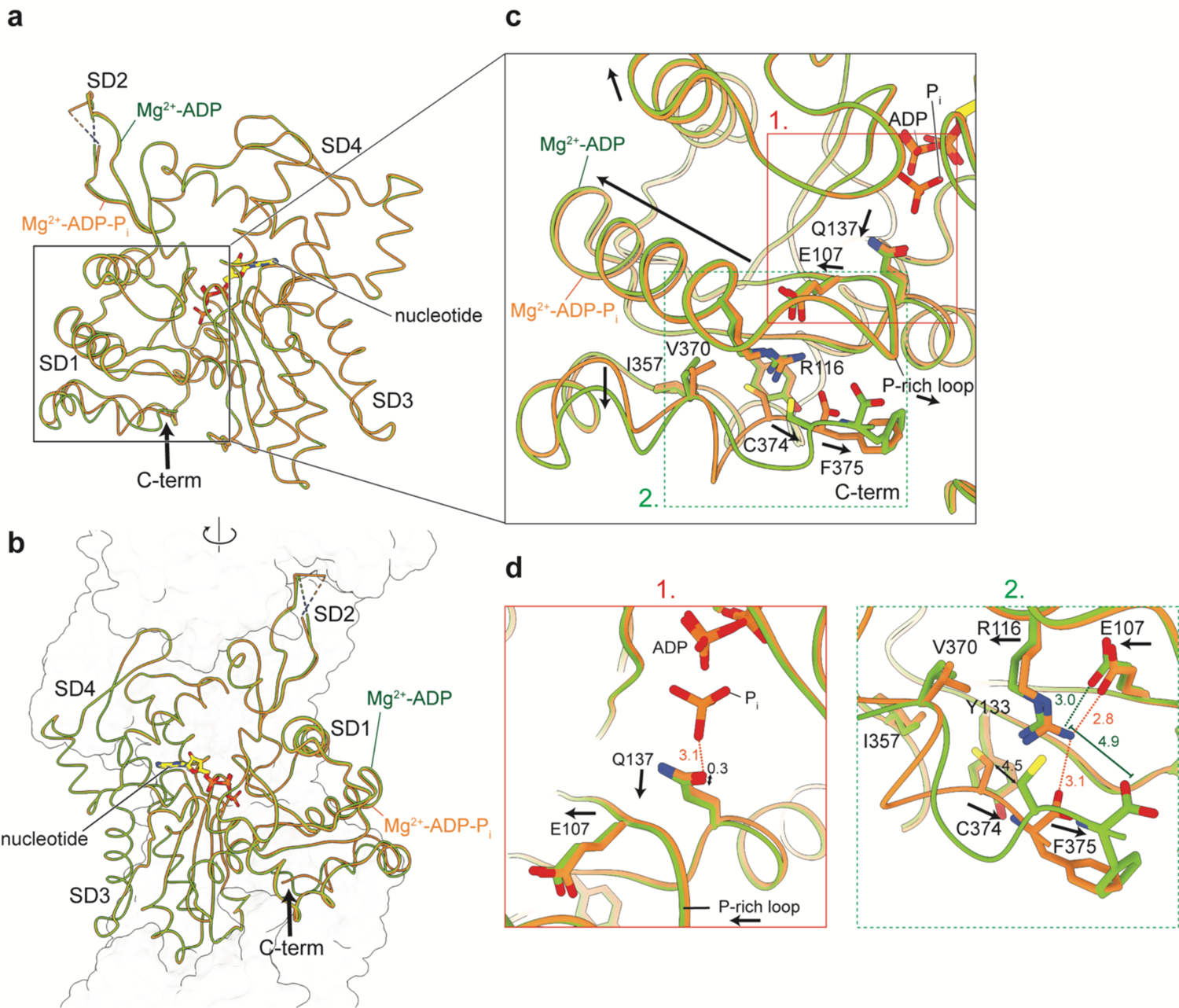
Structural coupling of the nucleotide binding site to the filament exterior after P_i_ release. **a, b** Overlay of one actin subunit in the Mg^2+^-ADP-P_i_ and Mg^2+^-ADP structures with annotated subdomains, shown in two orientations. The location of the C-terminus (C-term) is accentuated with a large arrow. In **(b)**, the surface-contour of other actin subunits within the filament is depicted. **c** Differences in the SD1 of F-actin in the Mg^2+^-ADP-P_i_ and Mg^2+^-ADP structures. Residues thought to be important for the movement are annotated. **d** Zoom of the nucleotide binding site (1.) and C-terminal region (2.) of the SD1. In (**c)** and **(d)**, arrows depict the direction of the putative movement from the Mg^2+^-ADP-P_i_ to the Mg^2+^-ADP structure. All distances are shown in Å. Distances shown in the Mg^2+^-ADP-P_i_ structure are colored orange, whereas those in the Mg^2+^-ADP structure are colored green.

**Supplementary Fig. 16.**
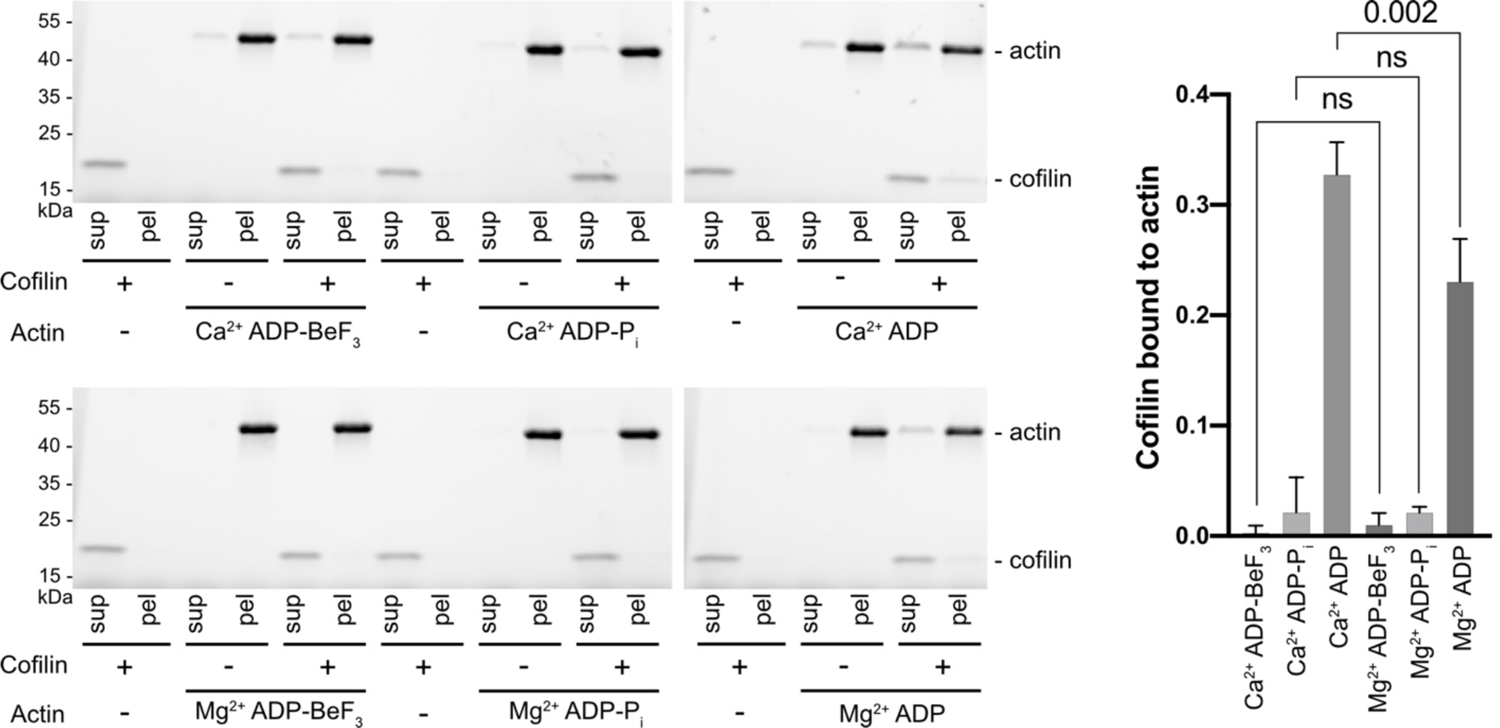
Cofilin co-sedimentation assays. Representative stain-free SDS PAGE gel images of co-sedimentations of cofilin with Ca^2+^-F-actin and Mg^2+^-F-actin in ADP-BeF_3_^-^, ADP-P_i_ and ADP states. The graph depicts the fraction of cofilin in the F-actin pellet from 3 independent assays. The data are presented as mean values, statistical analysis was performed by ordinary one-way ANOVA test. Raw gel images are provided in Supplementary Fig. 17.

**Supplementary Fig. 17.**
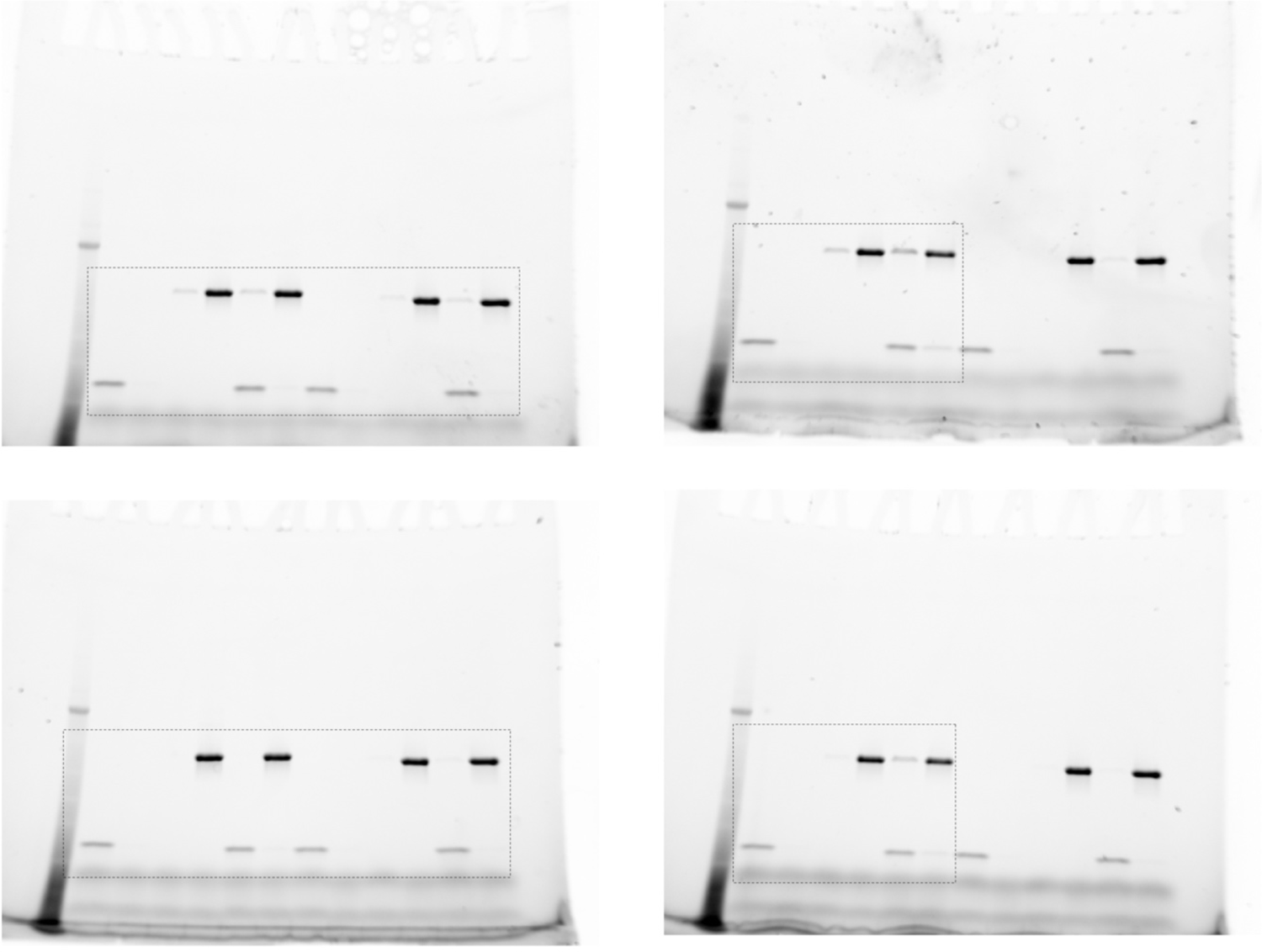
Uncropped SDS PAGE gels of cofilin co-sedimentation assays. The figure shows the complete gel images from Supplementary Fig. 16, in the same order. The rectangle shows the portion of the gel that was depicted in Supplementary Fig. 16.

**Supplementary Fig. 18.**
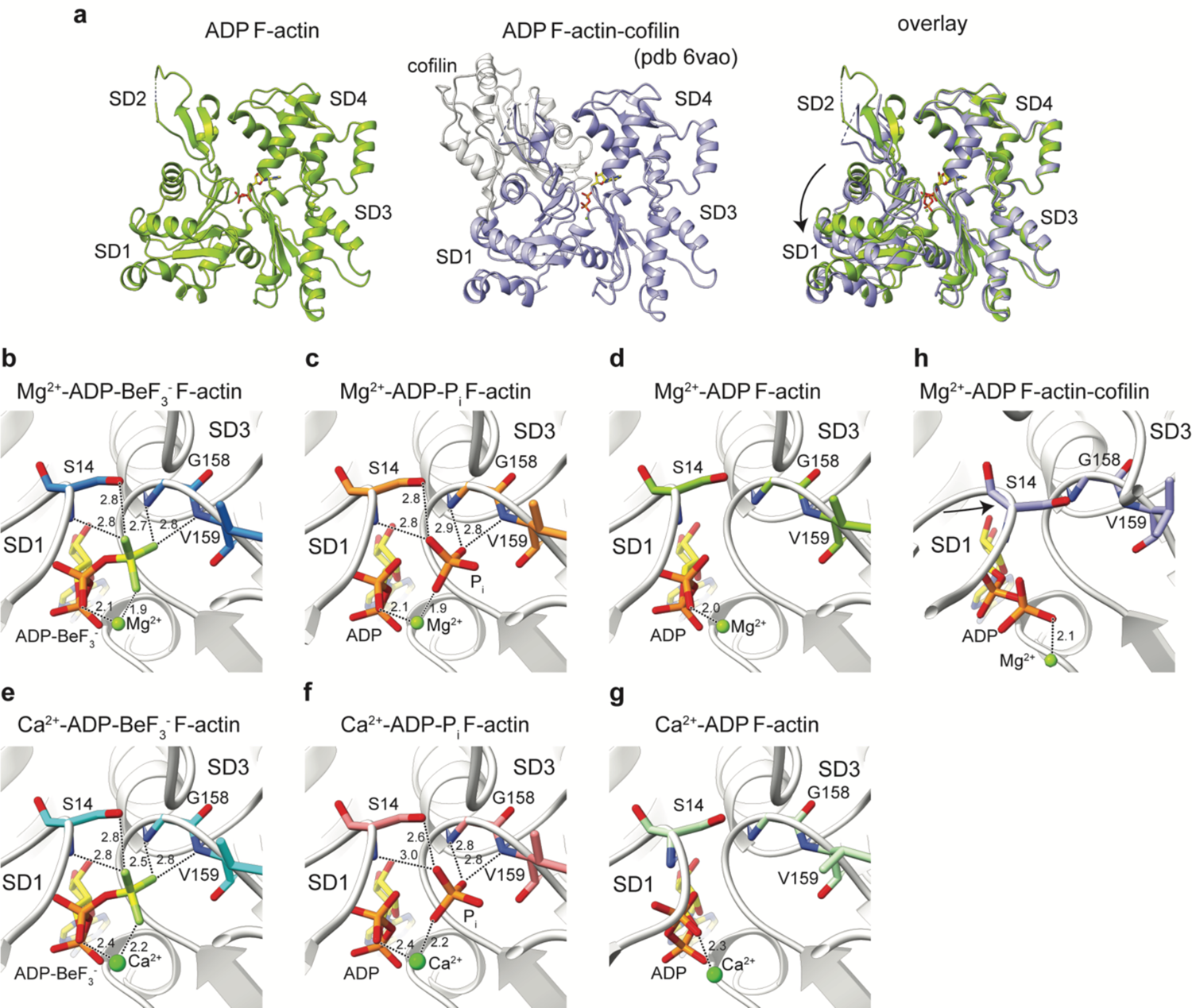
Cofilin-induced conformational changes in F-actin. **a** Structure of a single subunit of ADP F-actin (left panel) and cofilin-decorated ADP F-actin (middle panel). The right panel depicts an overlay between the two structures; cofilin is hidden for clarity. The arrow indicates the SD1 and SD2 rotation in F-actin upon cofilin binding. **b-h** Arrangement of the SD1 and SD3 at the nucleotide binding sites in Mg^2+^-ADP-BeF_3_^-^ **(b)**, Mg^2+^-ADP-P_i_ **(c)**, Mg^2+^-ADP **(d)**, Ca^2+^-ADP-BeF_3_^-^ **(e)**, Ca^2+^-ADP-P_i_ **(f)**, Ca^2+^-ADP **(g)** and cofilin-decorated Mg^2+^-ADP **(h)** structures of F-actin. In panel **h**, the arrow depicts the cofilin—induced SD1 movement.

